# KDM6 demethylases mediate EWSR1-FLI1-driven oncogenic transformation in Ewing Sarcoma

**DOI:** 10.1101/2023.01.24.524910

**Authors:** Elisabet Figuerola-Bou, Carla Rios-Astorch, Enrique Blanco, María Sánchez-Jiménez, Pablo Táboas, Guerau Fernández, Soledad Gómez, Oscar Muñoz, Pol Castellano-Escuder, Sara Pérez-Jaume, Estela Prada, Silvia Mateo-Lozano, Nicolo Riggi, Alexandra Avgustinova, Cinzia Lavarino, Luciano Di Croce, Sara Sánchez-Molina, Jaume Mora

**Affiliations:** Pediatric Cancer; Institut de Recerca Sant Joan de Deu; Esplugues de Llobregat (Barcelona), 08950; Spain; Center for Genomic Regulation (CRG); Barcelona Institute of Science and Technology; Barcelona, 08003; Spain; Universitat Pompeu Fabra (UPF); Barcelona, 08002; Spain; Molecular Genetics Department; Institut de Recerca Sant Joan de Déu; Esplugues de Llobregat (Barcelona), 08950; Spain; Biomarkers and Nutritional & Food Metabolomics Research Group; Department of Nutrition, Food Science and Gastronomy, University of Barcelona; Barcelona, 08028; Spain; Institute of Pathology, Centre Hospitalier Universitaire Vaudois; Faculty of Biology and Medicine, University of Lausanne; Lausanne, 1004; Switzerland; Pediatric Cancer Center Barcelona (PCCB); Hospital Sant Joan de Déu; Esplugues de Llobregat (Barcelona), 08950; Spain; Institucio Catalana de Recerca i Estudis Avançats (ICREA); Barcelona, 08010; Spain

**Keywords:** Ewing sarcoma, EWSR1-FLI1, KDM6A, KDM6B, enhancer, H3K27me3

## Abstract

Ewing Sarcoma (EwS) is an aggressive bone and soft tissue tumor driven by the fusion oncoprotein EWSR1-FLI1. This aberrant transcription factor binds to GGAA microsatellites, causing epigenetic reprogramming through the formation of active neo-enhancers in a permissive cellular context. Inhibition of the oncogene remains challenging and current efforts instead seek to exploit emergent epigenetic treatments targeting EWSR1-FLI1 cofactors. Here, stemming from the genome-wide redistribution of H3K27me3 upon expression of EWSR1-FLI1 in pediatric hMSC, we unravel the contribution of the H3K27me3 demethylases KDM6A and KDM6B in transcriptional activation at EWSR1-FLI1 enhancers. We found that KDM6A has a demethylase-independent role in recruiting the SWI/SNF member BRG1 at EWSR1-FLI1-primed enhancers containing single GGAA motif, which is critical for EwS tumor growth. Conversely, KDM6B demethylates H3K27me3 at EWSR1-FLI1-active enhancers containing multimeric GGAA repeats and its deletion synergizes with EZH2 inhibitors. Our results highlight KDM6 demethylases as EWSR1-FLI1 cofactors with potential for future targeted therapies.

## INTRODUCTION

Ewing Sarcoma (EwS) is a deadly neoplasm that affects the bone and soft tissues of children, adolescents and young adults (Grünewald et al., 2018). Whole-genome sequencing studies of EwS reported a remarkably genome stability with very low mutational burden (Brohl et al., 2014; Crompton et al., 2014). Like many developmental tumors EwS presented a maturation block during development conferred by the expression of an oncogenic driver (Behjati et al., 2021). The characteristic reciprocal chromosomal translocation most commonly involves the *EWSR1 RNA-binding protein 1* (*EWSR1*) gene in chromosome 11 and the *ETS transcription factor family member Fli-1 proto-oncogene* (*FLI1*) in chromosome 22 (Delattre et al., 1992). The resulting fusion protein retains the transactivation domain of *EWSR1* fused to the DNA-binding domain of *FLI1* and acts as an aberrant transcription factor (Lessnick et al., 1995; May et al., 1993). However, EWSR1-FLI1 can only achieve oncogenic transformation in the right cellular background (Patel et al., 2012; Riggi et al., 2008; von Levetzow et al., 2011). While EWSR1-FLI1 induces growth arrest or apoptosis in differentiated primary cells (Deneen and Denny, 2001; Lessnick et al., 2002), its expression in human mesenchymal stem cells (hMSC) recapitulates the EwS gene signature (Riggi et al., 2010; Riggi et al., 2008). Nevertheless, stronger induction of oncogenic targets in pediatric MSC (hpMSC) as opposed to adult origin highlights the need for a right permissive molecular framework (Riggi et al., 2010).

EWSR1-FLI1 exhibits affinity to bind DNA through GGAA motifs, a class of ETS specific response element, activating or repressing their targets based upon the number of motif repeats (Gangwal et al., 2010; Gangwal et al., 2008). At GGAA repeats EWSR1-FLI1 multimers establish *de novo* active enhancers (neo-enhancers) by promoting chromatin opening through recruitment of chromatin-modifying complexes. Indeed, in EwS cell lines and primary tumors a vast majority of EWSR1-FLI1 binding sites are decorated by H3K27ac, a post-translational histone mark associated to active enhancers (Riggi et al., 2014; Tomazou et al., 2015). Consistently, EWSR1-FLI1 exhibits scaffolding properties and recruits core members of the transcriptional activator SWItch/Sucrose Non-Fermentable (SWI/SNF) chromatin complexes, such as BAF155, at GGAA repeats that are critical for transcriptional activation (Boulay et al., 2017). Recently, we have described that RING1B, a member of the Polycomb group (PcG) family of proteins (Schuettengruber et al., 2017), facilitates EWSR1-FLI1 recruitment towards key enhancers (Sánchez-Molina et al., 2020). Although the mechanism behind EWSR1-FLI1 gene repression is less well understood, it has been proposed that binding of EWSR1-FLI1 monomers on single instances of the GGAA motif cause displacement of endogenous ETS transcription factors decreasing transcriptional activation (Riggi et al., 2014). Besides, a large number of transcriptional repressors such as *NKX2-2* are induced by EWSR1-FLI1, highlighting indirect mechanisms to mediate gene silencing (Owen et al., 2008).

The post-translational histone mark trimethylation of H3K27 (H3K27me3) decorates promoters and enhancers of repressed and bivalent genes (Ezponda and Licht, 2014). H3K27me3 mark restricts cell fate by limiting chromatin accessibility to key developmental genes, thus proper addition or removal of this mark in a timely manner is critical during development (Margueron and Reinberg, 2011). EZH2, the core member of the Polycomb Repressive Complex 2 (PRC2), is the primary enzymatic writer of H3K27me3 that maintain silencing patterning fundamental for cell identity and proper differentiation (Comet et al., 2016; Chammas et al., 2020). Removal of H3K27me3 concerted with promoter activation is mediated by members of the KDM6 family of demethylases, including KDM6A (UTX) and KDM6B (JMJD3), through its JmjC domain (Agger et al., 2007; Lee et al., 2007) and determine specification of human neural progenitor cells (Shan et al., 2020). KDM6A has been described as a partner of the Set1/MLL complex which is responsible for active gene expression by methylation of H3K4 at active enhancers (Lee et al., 2012; Wang et al., 2017), while KDM6B has been reported to cooperate with the transcription factor KLF4 in enhancer-driven reprogramming and with SMAD3 in neural specific enhancers (Fueyo et al., 2018; Huang et al., 2020).

Deregulation of the H3K27me3 balance that is governed by the coordinated enzymatic activity of EZH2 and KDM6A/KDM6B leads to differentiation defects and cancer (Das and Taube, 2020). EZH2 has been involved in the tumor progression mechanisms of a variety of cancer types (Kim and Roberts, 2016), while KDM6A and KDM6B have been identified in numerous malignancies with either oncogenic or tumor suppressor roles (Arcipowski et al., 2016; Benyoucef et al., 2016; Leng et al., 2020; Ntziachristos et al., 2014; van der Meulen et al., 2015). We previously demonstrated that EWSR1-FLI1 occupies weak repressed Polycomb chromatin states in hMSC (Sánchez-Molina et al., 2020). Although EWSR1-FLI1-bound GGAA repeats are devoid of H3K27me3 both in EwS cell lines and primary tumors (Riggi et al., 2014; Tomazou et al., 2015), in Human Umbilical Vein Endothelial Cells (HUVECs) and hpMSC these regions are extensively decorated with H3K27me3 (Boulay et al., 2018; Patel et al., 2012). EZH2 is a well-known directly activated target by EWSR1-FLI1 that blocks neuroectodermal and endothelial differentiation in EwS (Richter et al., 2009; Riggi et al., 2008). However, the precise mechanism behind H3K27me3 deposition and erasure equilibrium and its impact in EwS tumor development has not been addressed.

Here, we provide novel insights into H3K27me3 KDM6A/B removal dynamics in the context of EwS. First, we show H3K27me3 is redistributed genome-wide during EwS tumorigenesis in hpMSCs. Next, we demonstrate that KDM6A and KDM6B demethylases bind to the same genomic regions as EWSR1-FLI1, with KDM6A decorating EWSR1-FLI1 intermediate enhancers containing GGAA single motifs and KDM6B demethylases characteristically enriched at active enhancers containing multimeric GGAA. Importantly, both KDM6A and KDM6B demethylases are involved in EWS-related transcriptional activation in a demethylase-independent and dependent manner, respectively. Besides, KDM6A deletion delays tumor growth in xenografts and KDM6B perturbation synergize with specific inhibitors of EZH2. Our findings provide deep knowledge regarding specific functions of H3K27me3 KDM6 demethylases in EwS and new avenues for its treatment through the reversibility of epigenetic processes involved with H3K27me3 equilibrium.

## RESULTS

### H3K27me3 genome-wide redistribution upon EWSR1-FLI1 overexpression in hpMSCs

Several groups have documented a *bona fide* list of GGAA repeats bound by EWSR1-FLI1 to activate transcription (Bilke et al., 2013; Gangwal et al., 2008; Riggi et al., 2014). Recently, our group and others have demonstrated that overexpression of the oncogene triggers a loss of the H3K27me3 at promoters of genes that become transcriptionally activated in HUVEC cells and neural crest stem cells (NCSC) (Svoboda et al., 2014) (Sánchez-Molina et al., 2020). In an attempt to determine the global extension of such a decrease in H3K27me3, we analyzed published ChIP-seq data on H3K27me3 in hpMSC overexpressing the fusion oncogene (Boulay et al., 2018). We first examined the distribution of H3K27me3 in genes decorated by this mark in the hpMSC control condition (5 kb around TSS) and revealed a maximal decrease at 2 kb upstream the TSS upon EWSR1-FLI1 overexpression, concomitant with an increase in H3K27ac specific for TSS (Figure S1A). Nevertheless, Western blot analysis reported similar H3K27me3 levels in both conditions (Figure 1A), suggesting an overall genome-wide redistribution following the introduction of EWSR1-FLI1. In order to confirm this hypothesis, we segmented the genome into 3,069,655 bins of 1 kb and determined that averaged H3K27me3 signal strength along bins was highly correlated between control and EWSR1-FLI1 samples (Figure S1B). Of note, when highlighting with colors the bins of the genome containing H3K27me3 peaks, we observed a progressive loss of this mark in parallel with a gain of EWSR1-FLI1 signal upon expression of the oncogene. Conversely, EWSR1-FLI1 binding sites are originally decorated with H3K27me3 in the control condition (Figure 1B). As previously proposed (Sánchez-Molina et al., 2020), most genomic regions presenting H3K27me3 changes upon EWSR1-FLI1 expression are significantly enriched over regulatory sequences of bivalent genes in hpMSC (Figure S1C). Specifically, we identified 103,766 bins that presented a significant reduction of H3K27me3 upon EWSR1-FLI1 introduction (Down bins) (Figures S1B and S1D), corresponding to 2,879 genes involved in neural processes and metabolism (Figure S1E and Table S1). For instance, *NKX2-2*, a well-known EWSR1-FLI1 target, exhibited a gain of H3K27ac with a concomitant moderate H3K27me3 decay along its promoter and regulating enhancers (Figure 1C). We also identified 110,845 bins (2,621 genes) in the genome showing a significant gain of H3K27me3 upon EWSR1-FLI1 overexpression (Up bins) (Figures S1B and 1D). Functional analysis of such genes reported association with the TGF-beta receptor signaling pathway (Figure S1E and Table S1), in agreement with the well-known repressed state of genes from this pathway in EwS (Hahm et al., 1999). For instance, *TGFBI* was one of the genes in which a gain of H3K27me3 after EWSR1-FLI1 introduction was observed along the gene body (Figure 1D). Altogether, these results demonstrate that EWSR1-FLI1 targets regions decorated with H3K27me3 producing a redistribution of H3K27me3.

**Figure 1.**
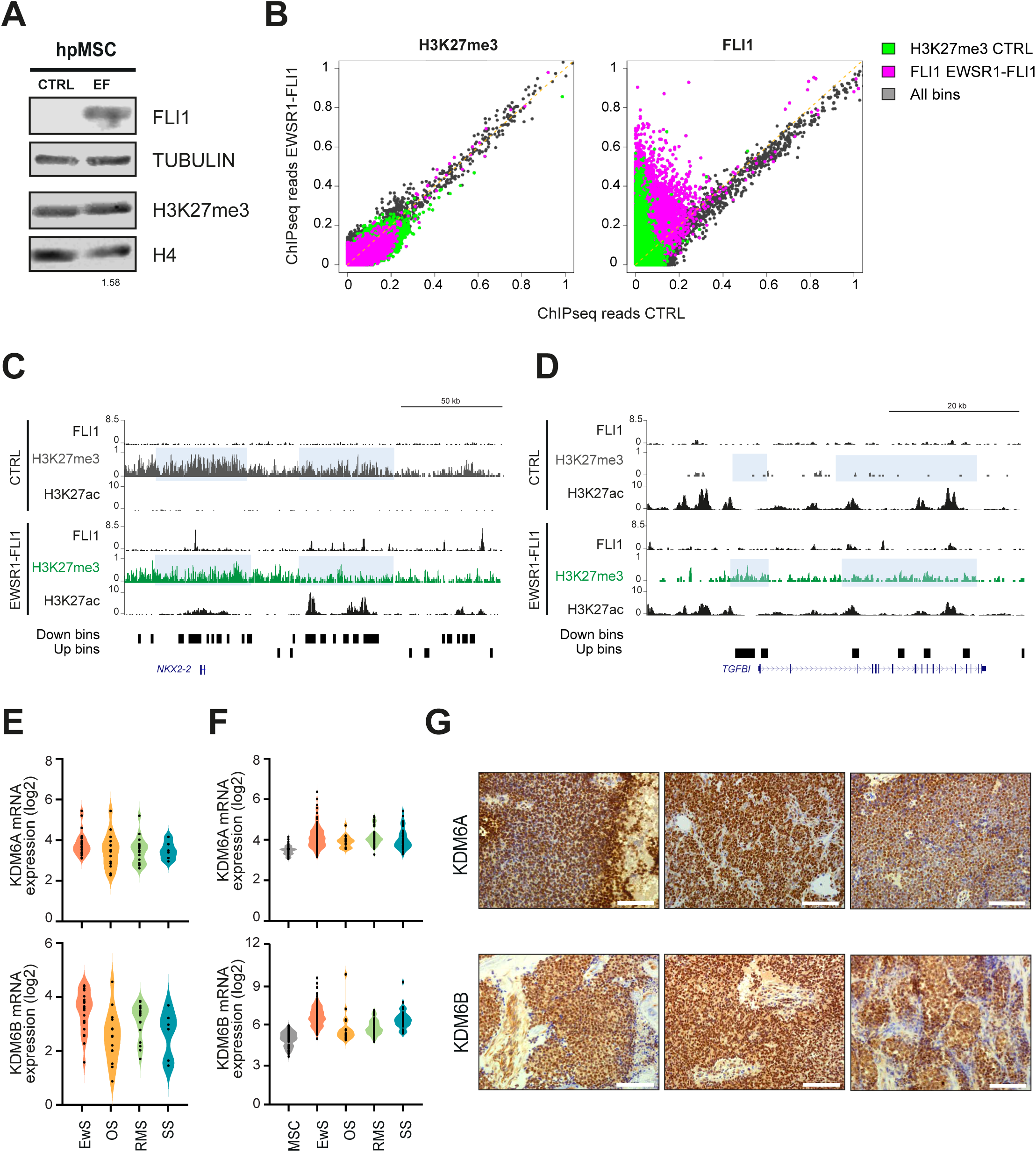
H3K27me3 genome-wide redistribution upon EWSR1-FLI1 overexpression in hpMSCs. (A) Western blot showing levels of FLI1 and H3K27me3 in whole cell and histone extracts, respectively, in control (CTRL) and upon EWSR1-FLI1 overexpression in hpMSC (EF). Tubulin and histone H4 were used as loading controls. Numbers below represent band quantification of H3K27me3 normalized to H4 and relative to CTRL. (B) Scatter plot depicting H3K27me3 (left) and FLI1 (right) ChIP-seq signal in 3,065,655 1 kb bins in CTRL (x-axis) and EWSR1-FLI1 (y-axis) hpMSC (R^2^=0.582, slope 0.703; and R^2^=0.074, slope 0.347, respectively). Bins containing ChIPseq peaks of H3K27me3 in CTRL hpMSC are highlighted in green, while bins containing FLI1 after EWSR1-FLI1 overexpression are shown in violet. (C) UCSC genome browser signal tracks for H3K27me3, H3K27ac, and FLI1 in CTRL and EWSR1-FLI1 hpMSC at the *NKX2-2* gene. Up or Down bins correspond to regions that gain or loss H3K27me3 upon EWSR1-FLI1 introduction, respectively, and are represented as black rectangles below tracks. Clusters of bins are highlighted in light blue. (D) Same as (C) at the *TGFBI* enhancer. (E) Violin plots representing mRNA levels of KDM6A (above) and KDM6B (below) in a panel of EwS, osteosarcoma (OS), rhabdomyosarcoma (RMS), and synovial sarcoma (SS) cell lines extracted from Barretina et al. (2012). (F) Same as (E) in primary sarcoma tumors from NCBI GEO public repository. MSCs derived from the healthy bone marrow were used as control tissue. (G) Immunohistochemical staining of KDM6A (above) and KDM6B (below) on sections of three representative primary EwS tumors from the cohort of 43 tumors from our institution counterstained with hematoxylin-eosin. White bar indicates 100µm scale. See also Figure S1 and Table S1.

Next, we hypothesized that KDM6A and KDM6B might be implicated in the prominent loss of H3K27me3 at promoters and enhancers of EWSR1-FLI1 target genes. Indeed, when evaluating the expression of KDM6A and KDM6B demethylases in a panel of cancer cell lines (Barretina et al., 2012), we found that while KDM6A expression was similar across tumor types, KDM6B was more expressed in EwS cell lines than in other sarcomas (Figure 1E). Similar trends were reported in 184 EwS tumors at diagnosis (GSE17679, GSE34620 and GSE37371 accessions), where we confirmed expression of both KDM6A and KDM6B consistently higher than MSC, while KDM6B expression in EwS tumors was particularly high compared to other sarcomas (Figure 1F). Analysis of KDM6A and KDM6B by immunohistochemistry in 43 EwS primary tumor specimens from newly diagnosed patients from our institution revealed that both demethylases were highly expressed by semi-quantitative histoscore (H-score) analysis (Figures 1G and S1F). No significant correlation between KDM6A and KDM6B expression in EwS tumors was found according to the statistical analysis of their H-score (Figure S1G). Taken together, RNA expression data and immunohistochemical studies indicate that KDM6A and KDM6B are highly expressed in EwS cell lines and in primary tumor samples.

### KDM6A and KDM6B co-localize genome-wide with EWSR1-FLI1 at primed and active enhancers

To shed light whether KDM6A and KDM6B demethylases are involved in H3K27me3 to H3K27ac switches in enhancers regulated by EWSR1-FLI1, we carried out chromatin immunoprecipitation followed by high-throughput sequencing (ChIP-seq) for KDM6A, KDM6B and EWSR1-FLI1 in the EwS cell line A673. We identified 3,737 peaks for KDM6A, 2,687 for KDM6B, and 4,800 for EWSR1-FLI1 (P value < 0.05 in all cases; FDR < 10^-3^ for KDM6A and EWSR1-FLI1, FDR < 10^-5^ for KDM6B) (Figure S2A and Table S2). At genomic level, we observed a clear preference for intergenic and genic regions in all cases (Figure 2A), which indicates a role at enhancers as previously described (Fueyo et al., 2018; Lee et al., 2012). KDM6B peaks were more enriched at promoter regions (10.1%), compared to EWSR1-FLI1 and KDM6A peaks. We next categorized KDM6A, KDM6B and EWSR1-FLI1 peaks into active, primed or poised enhancers, and in active or poised promoters based on Blanco et al. (2020). Primed enhancers are regulatory regions that correlate with the H3K4me1 mark and absence of H3K27ac (Drouin, 2014). Remarkably, KDM6A is more abundant at primed enhancers, whereas KDM6B mimics EWR1-FLI1 distribution and mainly associates to active enhancers (Figure 2B). In agreement with this, motif analysis found the characteristic single instance of the GGAA consensus on KDM6A sites, while KDM6B and EWR1-FLI1 peaks presented both forms (multiple and single) of the motif (Figure S2B). Gene association to peaks retrieves 1511 genes for KDM6A and 1207 genes for KDM6B, that were enriched in axon guidance, axonogenesis or nervous system development GO categories for both demethylases (Figure S2C and Table S3), indicating an important role of these demethylases in promoting neuronal differentiation as previously described (Shan et al., 2020). Primed enhancers define a state prior to activation that do not yield RNA and correlate with cell type specificity (Drouin, 2014; Maurya, 2021). Thus, the enrichment of KDM6A in primed enhancers supports the involvement of this enzyme in neural cell specification. Strikingly, when intersecting KDM6A and KDM6B peaks with the set of 697 EWSR1-FLI1-superenhancers defined by Tomazou et al. (2015), we found an overlap of 23% and 35%, respectively. To elucidate to which extend KDM6A and KDM6B co-localize with EWSR1-FLI1, we intersected peaks from the three entities and found a strong overlap of EWSR1-FLI1 with both demethylases (A-B-EF group) and with each demethylase separately (A-EF and B-EF) (Figure 2C). This result was validated in another EwS cell line, SK-ES-l, where we found 1163 common peaks between EWSR1-FLI1 and KDM6A/KDM6B (P value < 0.05; FDR < 10^-5^ in all cases) (Figure S2D). Motif analysis revealed a strong enrichment of GGAA multimeric repeats when EWSR1-FLI1 stands alone or together with KDM6B (B-EF group), while the presence of KDM6A is linked to the single GGAA motif (A-B-EF and A-EF groups), suggesting that KDM6A produces a unique set of EWSR1-FLI1 peaks that do not contain multimeric GGAA repeats but a single GGAA motif (Figure 2C). In agreement with our motif characterization, KDM6B together with EWSR1-FLI1 (B-EF group) strongly overlap with high levels of H3K27ac and H3K4me1 (i.e. active enhancers), while the presence of KDM6A inside a group is associated to lower levels of H3K27ac and enrichment in H3K4me1 (i.e. primed enhancers) (Figures 2D and 2E). Remarkably, KDM6B alone was found in a subset of transcriptionally active promoters (H3K4me3 and H3K27ac) (Figures 2D and S2E), suggesting differential functionalities for each demethylase. No significant enrichment on H3K27me3 was found in any collection of peaks (Figure S2E). We previously described RING1B co-localization genome-wide with EWSR1-FLI1 at active enhancers of key target genes (Sánchez-Molina et al., 2020). Interestingly, RING1B is enriched both in EWSR1-FLI1 enhancers containing KDM6A and KDM6B (B-EF, A-EF and A-B-EF group (Figure S2F). Our data support three classes of enhancers regarding their composition: (i) those where KDM6A and KDM6B co-localize with the oncogene at single GGAA motifs (e.g. *CDH11* gene, Figures 2F and S2H), (ii) KDM6A/EWSR1-FLI1 enhancers containing single GGAA motifs (e.g. *SMYD3*, Figure S2G); and (iii) those containing the overlap of KDM6B and EWSR1-FLI1 (without KDM6A) and enrichment at multimeric GGAA repeats (e.g. *NKX2-2* enhancer, Figures 2F and S2H). Altogether, the differential location and partnership identified for each of the KDM6 demethylases in EwS suggest that KDM6B specifies EWSR1-FLI1 for the most active regions and KDM6A signals for a specific set of enhancers that determines neural lineage.

**Figure 2.**
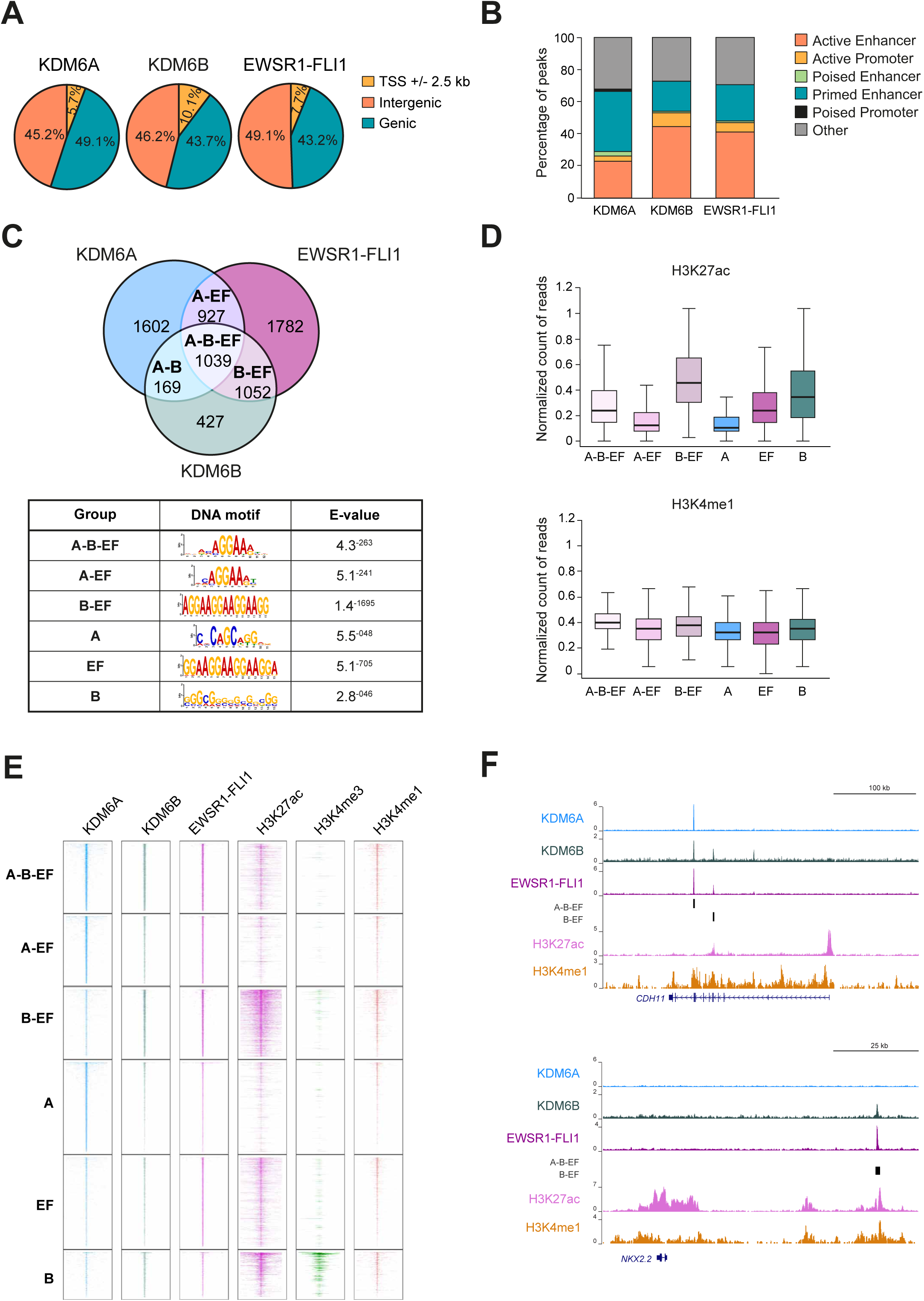
KDM6A and KDM6B co-localize genome-wide with EWSR1-FLI1 at primed and active enhancers. (A) Pie chart showing genomic distribution of KDM6A, KDM6B, and EWSR1-FLI1 peaks relative to functional categories including promoters (±2.5kb from TSS), genic regions (intragenic regions not overlapping with promoter) and intergenic regions (rest of the genome) in A673 cells. (B) Bar plot depicting percentage of total regulatory elements in the genome (active/poised/primed enhancers and active/poised promoters) overlapping KDM6A, KDM6B, and EWSR1-FLI1 peaks. (C) Venn diagram depicting the overlap of KDM6A, KDM6B, and EWSR1-FLI1 peaks in A673 cells. Table below shows top MEME DNA motifs and the corresponding E-value for every group of peaks. (D) Boxplot depicting the average ChIP-seq signal of H3K27ac (above) and H3K4me1 (below) on each subset of peaks. (E) Heatmap of KDM6A, KDM6B, EWSR1-FLI1, H3K27ac, H3K4me3, and H3K4me1 ChIP-seq signals for each group of peaks. (F) UCSC genome browser signal tracks for KDM6A, KDM6B, EWSR1-FLI1, H3K27ac, and H3K4me1 in A673 cells at the *CDH11* enhancer (above), and the *NKX2-2* enhancer (below). EWSR1-FLI1-KDM6B peaks with or without KDM6A (A-B-EF or B-EF, respectively) are represented as black bars below tracks. Error bars in (D) indicate SD. See also Figure S2 and Tables S2 and S3.

### Knockdown of KDM6A and KDM6B downregulates EWSR1-FLI1-activated targets

Our previous data point towards a different class of EWSR1-FLI1 enhancers when in partnership with KDM6A (primed enhancers) or with KDM6B (active enhancers). Indeed, the expression of genes in the proximity of enhancers containing binding sites of KDM6A is lower than those with KDM6B (Figure 3A). To gain insight into the role of both demethylases on EWSR1-FLI1-transactivation activity, we knocked-down KDM6A and KDM6B using a doxycycline-inducible system in EwS cell lines. We reported a significant knockdown of the demethylases at the protein level with two different shRNA sequences (KDM6A sh#1 and sh#2; KDM6B sh#1 and sh#2) in both A673 and SK-ES-1 cells (Figures 3B and S3A). In order to study genome-wide expression changes we carried out global transcriptome analysis by RNA-seq upon depletion with KDM6A (sh#1 and sh#2) and KDM6B (sh#1 and sh#2). Loss of each demethylase resulted in gene downregulation in A673 (P adj < 0.01 and log FC cut-off 0.5 and FPKM > 10, Figures 3C and S3B and Table S4) and SK-ES1 cell lines (P adj < 0.1 and log FC cut-off 0.32 and FPKM > 5, Figure S3C). Gene Set Enrichment Analysis (GSEA) in A673 cell line found that KDM6A and KDM6B are necessary for the activation of Epithelial-to-Mesenchymal Transition (EMT) genes (Figure 3D and Table S4). However, there is little overlap between both sets of deregulated genes upon demethylase knockdown (86 up-regulated and 34 down-regulated genes in common between KDM6A and KDM6B, data not shown), confirming different functionalities for each demethylase. All these results support a selective active role of KDM6A and KDM6B in the transcriptional activation network necessary for EwS tumorigenesis.

**Figure 3.**
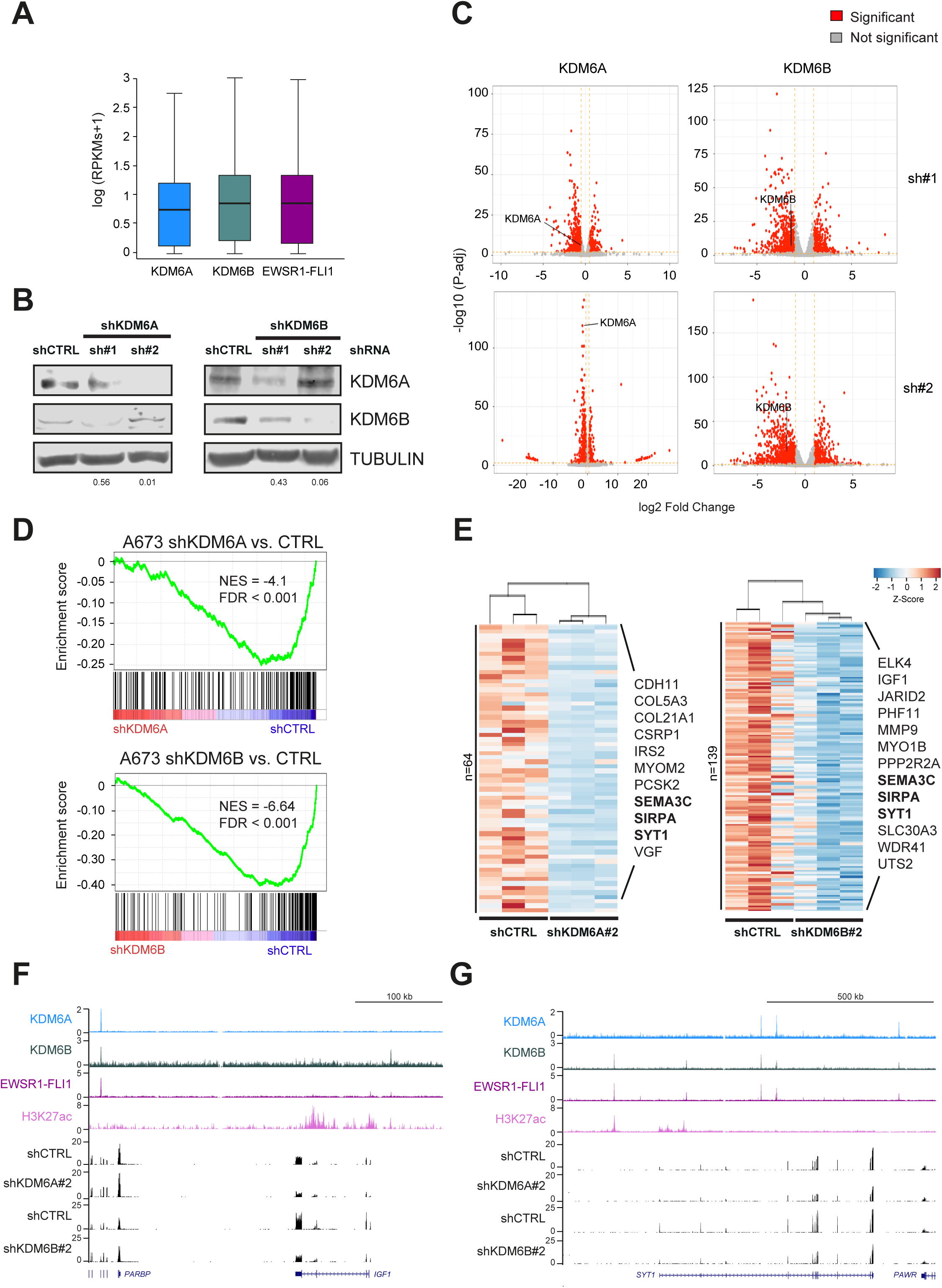
Knockdown of KDM6A and KDM6B downregulates EWSR1-FLI1-activated targets. (A) Boxplot representing the RNA-seq levels of KDM6A, KDM6B, and EWSR1-FLI1 target genes in A673 from Riggi et al. (2014). (B) Western blot showing levels of KDM6A and KDM6B in whole cell extracts upon KDM6A (left) or KDM6B (right) knockdown with two shRNA sequences (sh#1 and sh#2) at 72 hours in A673 cells. Tubulin was used as loading control. Numbers below represent band quantification of KDM6A or KDM6B normalized to tubulin and relative to shCTRL. (C) Volcano plots depicting log fold change (x-axis) and the associated log P-adjusted value (y-axis) of deregulated targets upon KDM6A (left) or KDM6B (right) knockdown with #sh1 or #sh2 sequences (above or below, respectively). Significant targets according to set cut-off are highlighted in red. (D) Gene Set Enrichment Analysis (GSEA) curves and Normalized Enrichment Scores (NES) for Hallmark collection of Epithelial to Mesenchymal Transition (EMT) in targets downregulated upon KDM6A (above) and KDM6B (below) knockdown in A673 cells. False Discovery Rate (FDR) is also shown. (E) Heatmap and dendrogram showing expression levels of genes in the vicinity of KDM6A-EWSR1-FLI1 and KDM6B-EWSR1-FLI1 ChIP-seq peaks (100 kb) that are significantly downregulated upon knockdown of each demethylase in A673 cells. n, indicates number of deregulated targets upon knockdown. Common KDM6A/KDM6B deregulated targets are in bold. (F) UCSC genome browser signal tracks for KDM6A, KDM6B, EWSR1-FLI1, and H3K27ac at *IGF1*. RNA-seq tracks for shKDM6A and shKDM6B (shRNA#2) and the corresponding shCTRL in A673 are shown. (G) Same as (**f**) at *SYT1*. Error bars in (A) indicate SD. See also Figure S3 and Tables S4 and S5.

To further understand the relevance of KDM6A and KDM6B in the transcriptional activation program of EWSR1-FLI1, we intersected genes associated to the genomic regions where these demethylases and EWSR1-FLI1 co-localize with deregulated targets in A673. To associate genes to peaks we captured every gene in a distance of 100 kb of both combinations of ChIP-seq peaks and obtained 3134 genes and 3326 genes in the vicinity of EWSR1-FLI1/KDM6A and EWSR1-FLI1/KDM6B, respectively. We then intersected these direct targets with the set of KDM6A and KDM6B significantly downregulated genes found upon knockdown and obtained 64 directly activated targets for KDM6A and 139 for KDM6B (Figure 3E and Table S5). GO analysis of such genes reported neural categories such as axonogenesis, as previously observed in Figure 2C, and cell migration (Figure S3D). Among the list of KDM6A/EWSR1-FLI1 activated targets we found *CDH11*, *IRS2* and *MYOM2* (Figures 3E and S3E); while for KDM6B/EWSR1-FLI1 we found targets such as *IGF1* and *JARID2* (Figures 3E, 3F and S3F). Moreover, certain genes (e.g. *SEMA3C* or *SYT1*) seem to be regulated by the combination of both KDM6A and KDM6B (A-B-EF) as described in Fig. 2 (Figures 3E and 3G).

### KDM6A recruits BRG1 to EWSR1-FLI1-activated enhancers in a demethylase-independent manner

To investigate in detail the demethylase role of KDM6A, we proceeded to study the effect of knocking out KDM6A using CRISPR/Cas9 genome editing. Our knockout completely abolished KDM6A expression in A673 cells without changes in EWSR1-FLI1 and KDM6B levels (Figure 4A). Despite loss of demethylase enzymatic activity, global levels of H3K27me3 as well as H3K4me1 and H3K27ac were not altered upon knockout in A673 and SK-ES1 cell lines (Figures 4B and S4A). We further validated by RT-qPCR the downregulation of genes containing KDM6A and EWSR1-FLI1 with KDM6B (A-B-EF group), while the expression of targets containing KDM6B and EWSR1-FLI1 (B-EF group) remained constant (Figure 4C).

**Figure 4.**
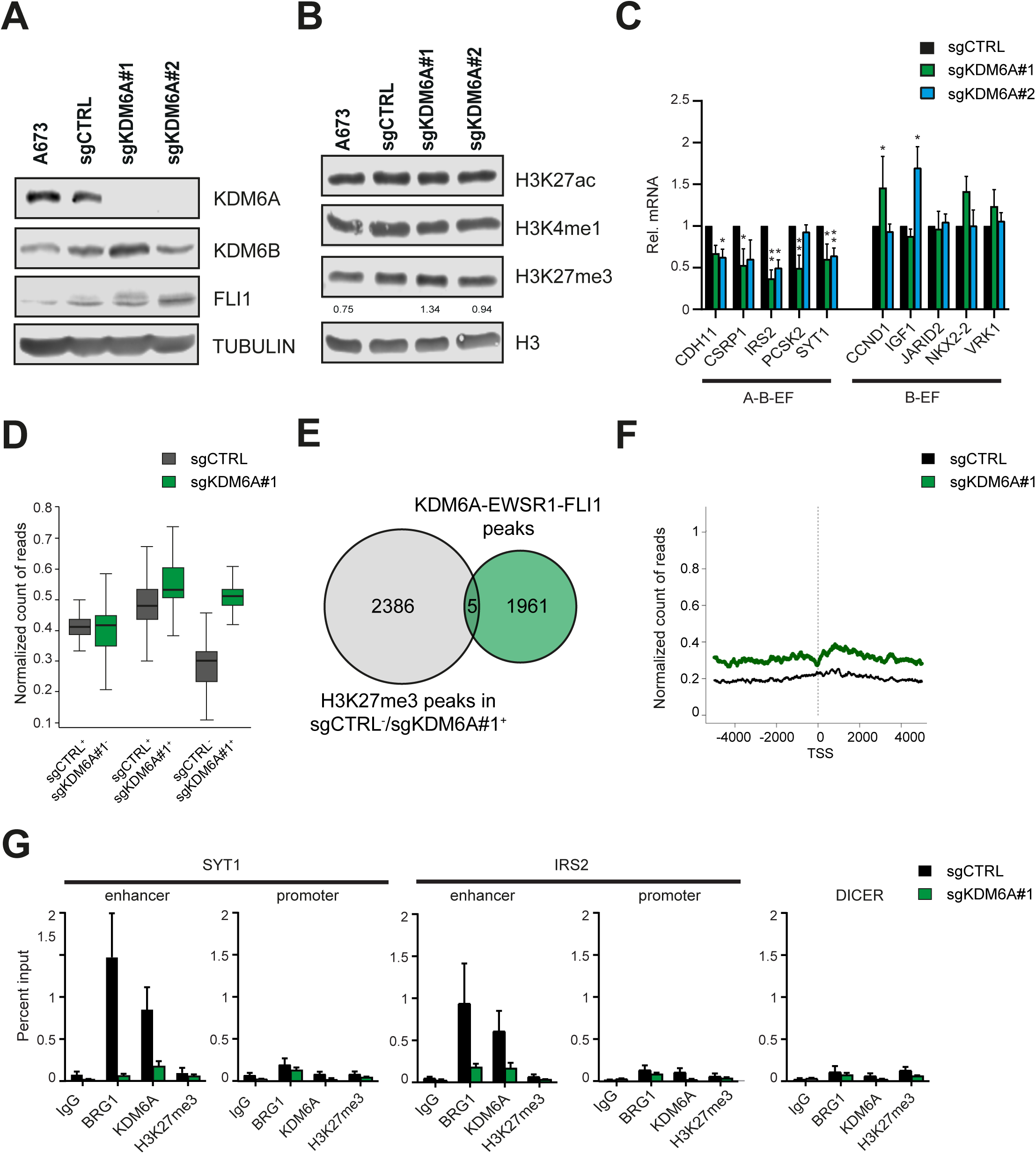
KDM6A recruits BRG1 to EWSR1-FLI1-activated enhancers in a demethylase-independent manner. (A) Western blot showing levels of KDM6A, KDM6B, and FLI1 in whole cell extracts upon KDM6A knockout with two sgRNA sequences (#1 and #2) in A673 cells. Tubulin was used as loading control. (B) Same as (A) with levels of H3K27ac, H3K4me1, and H3K27me3 in histone extracts upon KDM6A knockout in A673 cells. Histone H3 was used as loading control. Numbers below represent band quantification of H3K27me3 normalized to H3 and relative to non-targeting control (sgCTRL). (C) RT-qPCR determination of EWSR1-FLI1 targets with both KDM6A and KDM6B or with only KDM6B ChIP-seq peaks (A-B-EF and B-EF groups, respectively) in A673 knockout cells (sgRNA #1 and #2). *GAPDH* was used as housekeeping gene. (D) Boxplot depicting the average ChIP-seq signal of H3K27me3 in the three group of peaks in control (sgCTRL) and KDM6A knockout cells (sgKDM6A#1). 1278 peaks exclusive for sgCTRL or 1044 peaks for sgKDM6A#1 (sgCTRL^+^/sgKDM6A^-^ and sgCTRL^-^/sgKDM6A^+^, respectively), and 1347 peaks common in sgCTRL and sgKDM6A#1 (sgCTRL+/sgKDM6A^+^). (E) Venn diagram depicting the overlap between the 2391 ChIP-seq peaks of H3K27me3 in sgKDM6A#1 and the common set of 1966 peaks of KDM6A and EWSR1-FLI1. (F) Metagene plot showing H3K27me3 ChIP-seq signal of 341 KDM6A-activated targets from RNA-seq data at transcription start site (TSS) within 5000 kb window in sgCTRL and sgKDM6A#1. (G) ChIP-qPCR of BRG1, KDM6A, and H3K27me3 enrichment at the enhancer and promoter region of *SYT1* and *IRS2* KDM6A-activated targets upon KDM6A knockout with sgRNA#1. *DICER* was used as a negative control region. Statistical significance was determined by Kruskal-Wallis one-way analysis of variance (ANOVA) test (C). Error bars indicate SD (D). Error bars in (C) and (G) indicate SEM of three independent biological experiments; \*\**P* < 0.01, and \**P* < 0.05. See also Figure S4.

To inspect for the demethylase activity of KDM6A genome wide, we performed spike-in normalized ChIP-seq for H3K27me3 in A673 control and KDM6A KO conditions (sgCTRL and sgKDM6A A673 cells, respectively). At the genomic level, H3K27me3 distribution remains unchanged in the absence of KDM6A with strong enrichment in promoters (Figure S4B). To further investigate the effects of KDM6A KO on H3K27me3, we divided the resulting peaks into three classes: (i) commonly found in control and KDM6A KO, (ii) only found in the control condition, and (iii) only found upon KDM6A KO. Signal strength quantification of each class of peaks revealed an increase in H3K27me3 upon KDM6A KO in the set of peaks common for both conditions (sgCTRL+/sgKDM6A+) and in those new peaks that appeared upon demethylase deletion (sgCTRL-/sgKDM6A+, Figure 4D). Nevertheless, those new regions gaining H3K27me3 upon KDM6A KO were indirect effects of the depletion as its intersection with KDM6A peaks was very limited (Figure 4E). Indeed, the average profile of H3K27me3 around the TSS of 341 KDM6A-activated (downregulated upon KO) and 161 repressed targets (upregulated upon KO) revealed no correlation between changes on their gene expression and H3K27me3 levels (Figures 4F and S4C, respectively).

In order to accurately pinpoint the regions in the genome presenting higher H3K27me3 changes, we segmented the genome in bins of 1 kb and determined the average signal strength of the ChIP-seq of both conditions. This analysis retrieved 881,014 bins gaining (up) H3K27me3 and only 88,914 bins losing (down), with the set of up bins associated to 11658 genes and the down bins to 748, supporting a general trend towards H3K27me3 gain in sgKDM6A cells (sgCTRL-/sgKDM6A+; Figures S4D and S4E). Despite the global H3K27me3 increase upon KDM6A KO, only 36% of the 341 KDM6A-activated targets from RNA-seq data overlap with the 11,658 genes associated to up bins (Figures S4F). All these results suggest that KDM6A regulates expression of EWSR1-FLI1 targets in a demethylase-independent manner. KDM6A demethylase-independent functions have recently been proposed in the Kabuki syndrome, in the context of cell-induced differentiation with retinoic acid, mesoderm differentiation of embryonic stem cells and in lung cancer (Kim et al., 2014; Leng et al., 2020; Shpargel et al., 2017; Wang et al., 2012; Wang et al., 2017). To further confirm whether KDM6A acts without a demethylating activity, we evaluated levels of H3K27me3 on two KDM6A-activated target genes, *SYT1* and *IRS2*. Similar to previous results, H3K27me3 levels showed no significant changes by ChIP-qPCR upon KDM6A KO neither at the enhancer nor at the promoter regions (Figures 4G and S4G).

KDM6A has been described to physically associate with the SWI-SNF member BRG1/SMARCA4 (Gozdecka et al., 2018; Miller et al., 2010; Seunghee et al., 2012), which constitutes the central catalytic subunit that uses the energy derived from ATP-hydrolysis to remodel nucleosomes and regulate transcription (Phelan et al., 1999). To elucidate whether KDM6A was mediating transcriptional activation at enhancers through the recruitment of BRG1, we evaluated its enrichment in *SYT1* and *IRS2* enhancers. Interestingly, we observed that BRG1 was evicted from EWSR1-FLI1/KDM6A-bound enhancer regions following KDM6A deletion (Figures 4G and S4H). Altogether, these results suggest that KDM6A mediates recruitment of BRG1 at EWSR1-FLI1 enhancers facilitating transcriptional activation through a demethylase-independent mechanism.

### KDM6A is a critical factor for EwS engraftment and tumor growth

Based on the previous activity through EWSR1-FLI1 enhancers, we aimed to investigate the dependency from KDM6A in the tumor formation of EwS cells. First, to determine whether KDM6A affects EwS cell growth, we studied cell proliferation *in vitro* in KDM6A KO cells. We observed cell proliferation was significantly decreased on day 3 and day 6 for both sgKDM6A#1 and #2 knockout A673 cells (Figure 5A). Moreover, KDM6A-knockout cells exhibited a significant drop in their clonogenic capacity, compared to control and parental cells (Figures 5B and S5A). Remarkably, observed cell proliferation and transformation defects were not due to alterations on cell cycle (Figure 5B). Next, we subcutaneously injected sgCTRL and sgKDM6A #1 and #2 knockout A673 cells into nude athymic mice and monitored tumor growth. Xenografts of parental cells were also included as an additional control for tumor growth. Interestingly, we found KDM6A-knockout tumors (#1 and #2) showed a delay in tumor growth compared to control or parental-derived tumors (Figures 5C and S5C). At 17 days post-injection tumors were significantly smaller for sgKDM6A#1 (tumor volume mean of 121 and 162 mm^3^ for sgKDM6A#1 and #2, and 252.5 and 280mm^3^ for A673 and sgCTRL tumors, respectively) (Figure S5D). Moreover, median survival increased from 28 and 25 days for parental and control cells to 35 and 30 days for sgKDM6A#1 and #2, respectively (Figure 5D). Importantly, tumor engraftment was significantly perturbed in KDM6A knockout tumors with a median tumor engraftment of 20 and 17 days for sgKDM6A#1 and #2, respectively, against 16 days of sgCTRL and parental tumors (Figure 5E). We confirmed downregulation of KDM6A by RT-qPCR, immunohistochemistry and Western Blot, (Figures 5F, 5G and S5E). Importantly, we observed a decreased expression of enhancer-bound KDM6A/EWSR1-FLI1 targets, *CDH11* and *IRS2* (Figure 5F), supporting the transcriptional activating role of KDM6A at these oncogenic targets. Immunohistochemical analysis of tumors confirmed the expression of the EwS marker CD99 and lower expression of the proliferation marker Ki-67 in sgKDM6A-derived tumors, especially those from sgKDM6A#1 group (Figure 5G). Altogether, these results confirm KDM6A as a critical factor in EwS tumor growth.

**Figure 5.**
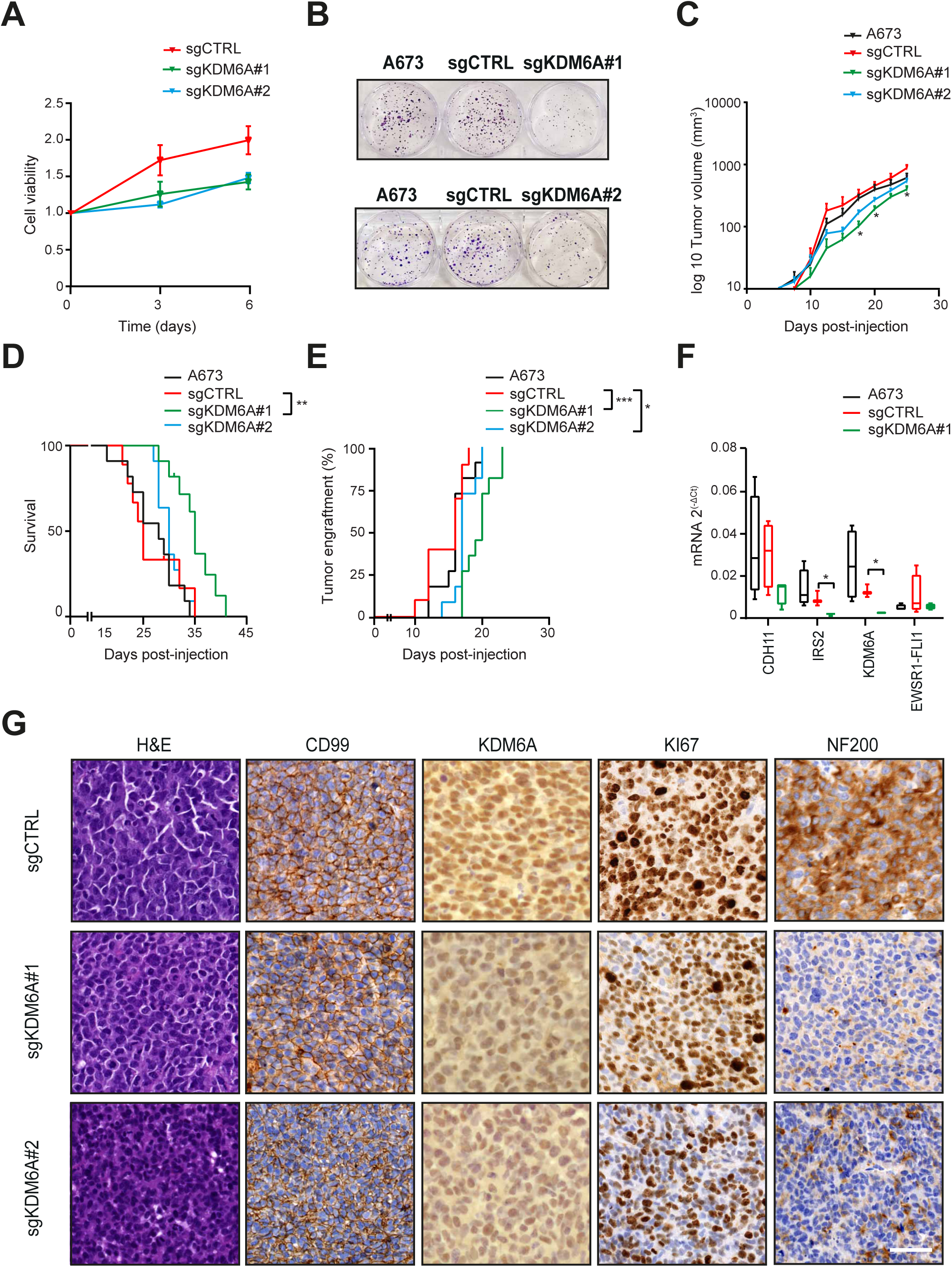
KDM6A is a critical factor for EwS engraftment and tumor growth. (A) Cell viability assay upon KDM6A knockout (sgRNA #1 and 2) in A673 cells compared to non-targeting control (sgCTRL) at 3 and 6 days. Values are relative to day 0. (B) Colony formation assay of KDM6A knockout cells (sgRNA#1 and 2) compared to parental cells and sgCTRL in A673. (C) Tumor growth curve of the average volume of xenografts derived from subcutaneous injection of KDM6A knockout cells (sgRNA#1 and 2) (n=11 each) compared to parental and sgCTRL A673 cells (n=11 and 9, respectively) in nude athymic mice from 0 to 24 days post-injection. (D) Kaplan-Meier representing the percentage of xenograft tumors that reach a tumor volume of 1000 mm^3^ within days post-injection in parental, sgCTRL and KDM6A knockout (sgRNA#1 and 2) in A673 cells. (E) Kaplan-Meier representing the percentage of xenograft tumors that reach a tumor volume of 150 mm^3^ within 24 days post-injection in parental, sgCTRL and KDM6A knockout (sgRNA#1 and 2) in A673 cells. Log-rank (Mantel Cox) test. (F) RT-qPCR determination of KDM6A targets *CDH11* and *IRS2*. *EWSR1-FLI1* and *KDM6A* are also shown. *GAPDH* was used as housekeeping gene. (G) Immunohistochemistry staining of CD99, KDM6A, Ki67, and NF200 on sections of tumors excised from sgCTRL and sgKDM6A (sgRNA#1 and 2) xenografts of A673 cells. CD99 was used as positive control for EwS cells, Ki67 for cell proliferation, and hematoxylin-eosin for histopathological evaluation of tissue. White scale bar represents 50 µm. Statistical significance was determined by two-way ANOVA test with Tukey post hoc correction (A), two-way ANOVA with Geiser-Greenhouse correction (C) and Kruskal-Wallis one-way ANOVA test (F). Survival analysis was performed by Log-rank (Mantel Cox) test (D, E). Error bars indicate SEM (A, C and F) of three independent biological experiments; \*\**P* < 0.01, and \**P* < 0.05. See also Figure S5.

Previously, we found KDM6A targets enriched in categories related to neural differentiation. Neurofilament (NF) proteins are cytoskeleton proteins that maintain neural axons and dendrites (Steinert and Roop, 1988). Neoplastic cells of neural origin or those exhibiting neural differentiation express NF. EwS tumors show positivity to cell surface antigens related to neuroectodermal lineage (Lipinski et al., 1986) and intracellular markers such as neuron specific enolase (NSE) and NF proteins (Cavazzana et al., 1987; Moll et al., 1987). The heavy chain of neurofilament protein NF200 is highly expressed in EwS cells and tumors (Lizard-Nacol et al., 1989; Lizard-Nacol et al., 1992). Considering that KDM6A targets developmental neural pathways in EwS, we hypothesized that neural markers such as NF200 could be perturbed in KDM6A-knockout tumors. Indeed, NF200 is highly expressed in publicly available datasets of EwS tumors and cell lines (Figure S5F and S5G). Furthermore, immunohistochemical analysis revealed lower levels of NF200 in KDM6A-knockout tumors as compared with those from control group (Figure 5G). Altogether, our data indicates KDM6A exerts its critical role on EwS tumor growth and engraftment likely regulating neural pathways *in vivo*.

### KDM6B knockout sensitizes EwS cells to the EZH2 inhibitor GSK126

KDM6A and KDM6B in conjunction with EWSR1-FLI1 participate in enhancers of neural genes with a different configuration of GGAA motifs, suggesting that both demethylases do not necessarily play the same function in EwS. In order to understand whether the role of KDM6B in EwS cells is associated to its demethylase activity in EwS cells, we investigated H3K27me3 dynamics in KDM6B depleted cells. First, we confirmed knockout of KDM6B in A673 cell lines, while levels of EWSR1-FLI1 and KDM6A remain constant (Figure 6A). Next, we evaluated the levels of H3K27me3 by Western blot in KDM6B-knockouts and knockdown. Similar to KDM6A (Figure 4B), we found that the depletion of the demethylase did not alter the H3K27me3 levels globally (Figures 6B and S6A). We confirmed by RT-qPCR that expression of targets potentially regulated by enhancers containing KDM6B and EWSR1-FLI1 (B-EF group) was exclusively dependent on KDM6B expression, while KDM6B knockout did not affect expression of targets containing enhancers with KDM6A binding sites (A-B-EF group, Figure 6C). Then, we inspected H3K27me3 in KDM6B and EWSR1-FLI1 targets by ChIP-qPCR and observed an H3K27me3 increase at these regions upon KDM6B knockout (Figures 6D and S6B). Moreover, when we treated EwS cells with the KDM6A/B demethylase inhibitor GSK-J4, we found that the expression of EWSR1-FLI1-activated targets was decreased strongly in those targets containing only KDM6B, suggesting GSK-J4 phenocopies expression changes induced upon KDM6B deletion (Figure S6C and S6D). The fact that KDM6B knockout led to a lower clonogenic capacity of EwS cells suggested that its demethylase activity in EWSR1-FLI1 targets might be essential (Figures 6E and S6E). Our results confirm therefore the demethylase-independent functions we observed for KDM6A and suggest that KDM6B exerts its functions as demethylase over EWSR1-FLI1 targets.

**Figure 6.**
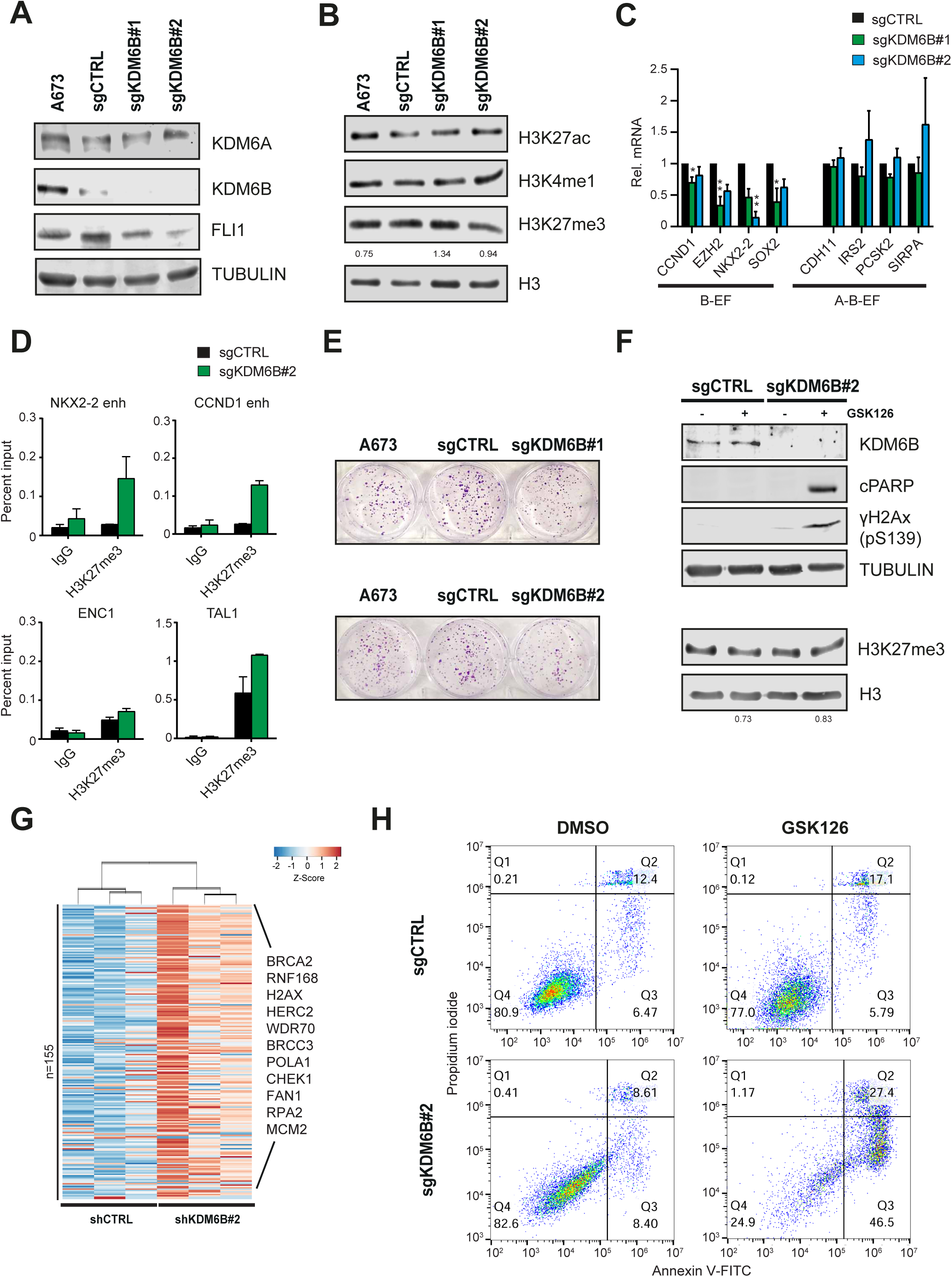
KDM6B knockout sensitizes EwS cells to the EZH2 inhibitor GSK126. (A) Western blot showing levels of KDM6A, KDM6B, and FLI1 in whole cell extracts upon KDM6B knockout with two sgRNA sequences (#1 and #2) in A673 cells. Tubulin was used as loading control. (B) Same as (A) with levels of H3K27ac, H3K4me1, and H3K27me3 in histone extracts upon KDM6B knockout in A673 cells. Histone H3 was used as loading control. Numbers below represent band quantification of H3K27me3 normalized to H3 and relative to non-targeting control (sgCTRL). (C) RT-qPCR determination of EWSR1-FLI1 targets with both KDM6A and KDM6B or with only KDM6B ChIP-seq peaks (A-B-EF and B-EF groups, respectively) in A673 knockout cells (sgKDM6B#1 and #2). *GAPDH* was used as housekeeping gene. (D) ChIP-qPCR of H3K27me3 enrichment at the enhancer region of *NKX2-2* and *CCND1* genes upon KDM6B knockout with sgRNA#2. *ENC1* and *TAL1* were used as negative and positive control regions, respectively. (E) Colony formation assay of KDM6B knockout cells (sgRNA#1 and 2) compared to parental cells and sgCTRL in A673. (F) Western blot showing levels of KDM6B, cleaved-PARP-1 (c-PARP), and phosphorylation of S139 in variant gamma-H2A.x (ɣ-H2Ax) in whole cell extracts (above) and H3K27me3 in histone extracts (below) from sgCTRL and sgKDM6B#2 A673 cells treated with vehicle or GSK126 inhibitor (15 µM for 24 hours). Tubulin and histone H3 were used as loading controls for whole cell or histone extracts, respectively. Numbers below represent band quantification of H3K27me3 normalized to H3 and relative to sgCTRL or sgKDM6B#2 treated with vehicle. (G) Heatmap and dendrogram showing expression of genes within the Hallmark collection of Double Strand Break Repair from GSEA analysis in shCTRL and shKDM6B#2, that are significantly downregulated upon KDM6B knockdown in A673 cells. n, indicates number of deregulated targets upon KDM6B knockdown. (H) Representative flow cytometry plots showing Annexin V-FITC/PI staining of sgCTRL and sgKDM6B#2 cells treated with vehicle or GSK126 inhibitor at 15 µM for 24 hours. Numbers in each quadrant indicate percentage of cells. Statistical significance was determined by Kruskal-Wallis one-way ANOVA test (C) and error bars indicate SEM (C and D) of three independent biological experiments; ***P < 0.001, and **P < 0.01. See also Figure S6 and Table S6.

Loss of function or inactivating mutations in KDM6A have been shown to sensitize to EZH2 inhibitors (Ezponda et al., 2017; Ler et al., 2017; Wu et al., 2018). Although EwS cells were reported to be resistant to EZH2 inhibition, the methyltransferase inhibitor GSK126 has been widely used to epigenetically de-repress key targets in EwS (Kailayangiri et al., 2019; Krook et al., 2016; Ryland et al., 2015). To further elucidate whether depletion of KDM6A/KDM6B demethylases would prime EwS cells to EZH2 inhibition we analyzed cell death in KDM6A/KDM6B knockout EwS cells upon GSK126 treatment. Only KDM6B knockout cells treated with the EZH2 inhibitor showed an increase in cleaved PARP-1 (c-PARP) and DNA damage marker ɣ-H2Ax levels by Western blot, while H3K27me3 decreased accordingly upon GSK126 treatment in KDM6A and KDM6B knockouts (Figures 6F and S6F). Consistently, we observed that depletion of KDM6B increased the expression of genes related to double-strand break repair pathways (Figure 6G and Table S6). Moreover, GSK126 exposure was sufficient to increase apoptosis by Annexin V staining in KDM6B KO cells, in contrast to KDM6A depletion where cells remained unaltered (Figures 6H and S6G). Furthermore, combination of GSK126 with GSKJ4 produce a synergistic response in apoptotic percentage (Figures S6H and S6I). These results suggest that EZH2 inhibition synergizes with the DNA damage infringed by KDM6B knockout in EwS cells.

## DISCUSSION

The importance of H3K27me3 redistribution in cancer has been previously reported in high-grade glioma, lymphoma, and melanoma, concomitant with a gain of function of EZH2 (Bender et al., 2013; Souroullas et al., 2016). In the present work, we performed an extensive genome-wide study of H3K27me3 redistribution upon EwS overexpression. Although the maintenance of the overall levels of H3K27me3 was previously described upon EWSR1-FLI1 knockdown, data on EZH2 and our results suggests gene specific changes are relevant for the tumorigenic process (Richter et al., 2009; Tomazou et al., 2015). Consistently, our findings indicate a genome-wide gain of H3K27me3 in genes from relevant EWSR1-FLI1 pathways, correlating with the important role of EZH2 in EwS tumorigenesis (Riggi et al., 2008). Furthermore, we confirmed a global loss of H3K27me3 at repressed regions in hpMSC once EWSR1-FLI1 is bound, as already proposed by Boulay et al. (2018). Altogether, our results refine the understanding of the early steps of EwS tumorigenesis and the role of H3K27me3 mark in defining the transformed epigenome of EwS.

We demonstrated that KDM6A and KDM6B promote the oncogenic process set by EWSR1-FLI1. Both demethylases reduce the clonogenic capacity of EwS and KDM6A depletion decreases not only tumor growth but tumor engraftment, suggesting an important role of KDM6A in early stages of tumorigenesis. We show KDM6A and KDM6B are cofactors of EWSR1-FLI1 and RING1B at distinct subsets of active enhancers in EwS. Moreover, their depletion alters pathways related to neural development and correlates with phenotypical expression of genes involved in EMT, which is intimately linked to the EwS metastatic processes (Pedersen et al., 2016). KDM6A has been identified as one of the 299 cancer driver genes in the cancer genome atlas project (TCGA) (Bailey et al., 2018). Its role in cancer development is cell-context specific, acting as a tumor suppressor or oncogenic factor. Loss-of-function mutations in KDM6A have been identified in cancers affecting males across multiple histiotypes, including T-ALL, pancreatic cancer, B-cell lymphoma, and medulloblastoma (Andricovich et al., 2018; Li et al., 2018; Robinson et al., 2012; van der Meulen et al., 2015). Interestingly, EwS is listed in the 12th position of cancer malignancies harboring higher mRNA expression of KDM6A (Barretina et al., 2012). Contrary to the dual role of KDM6A in cancer, KDM6B has been only correlated with cancer progression and a specific signature for EMT in lung and breast cancer (Lee et al., 2021; Ramadoss et al., 2012).

Most cancer studies deciphering the mechanisms behind KDM6A and KDM6B are based on independent approaches for each demethylase or its enzymatic inhibition by GSK-J4. Here, we performed a genome-wide analysis of both KDM6A and KDM6B in two EwS cell lines showing that both proteins co-localize with EWSR1-FLI1. Our study demonstrates that KDM6A and KDM6B collaborate with EWSR1-FLI1 through two different classes of enhancers that contribute distinctively to the oncogenic process: KDM6A enhancers, enriched in single instances of the GGAA motif, and KDM6B enhancers, which are characterized by multimeric GGAA repeats (Figure 7). Previous work by Riggi et al. (2014) described the single GGAA regions as EWSR1-FLI1-repressed elements where endogenous transcription factors had been displaced by the oncogene. Although these regions are devoid of the repressive mark H3K27me3, they are decorated with low levels of H3K27ac and transcription, correlating with our observation of KDM6A linked to primed enhancers. This type of enhancers constitute an intermediate state between active and repressed enhancers (Drouin, 2014; Maurya, 2021). Indeed, in our study KDM6A-associated genes containing GGAA single motifs are less transcribed than those enriched with KDM6B. Further analysis of the nature and implications of these primed enhancers and their differential regulation is needed.

**Figure 7.**
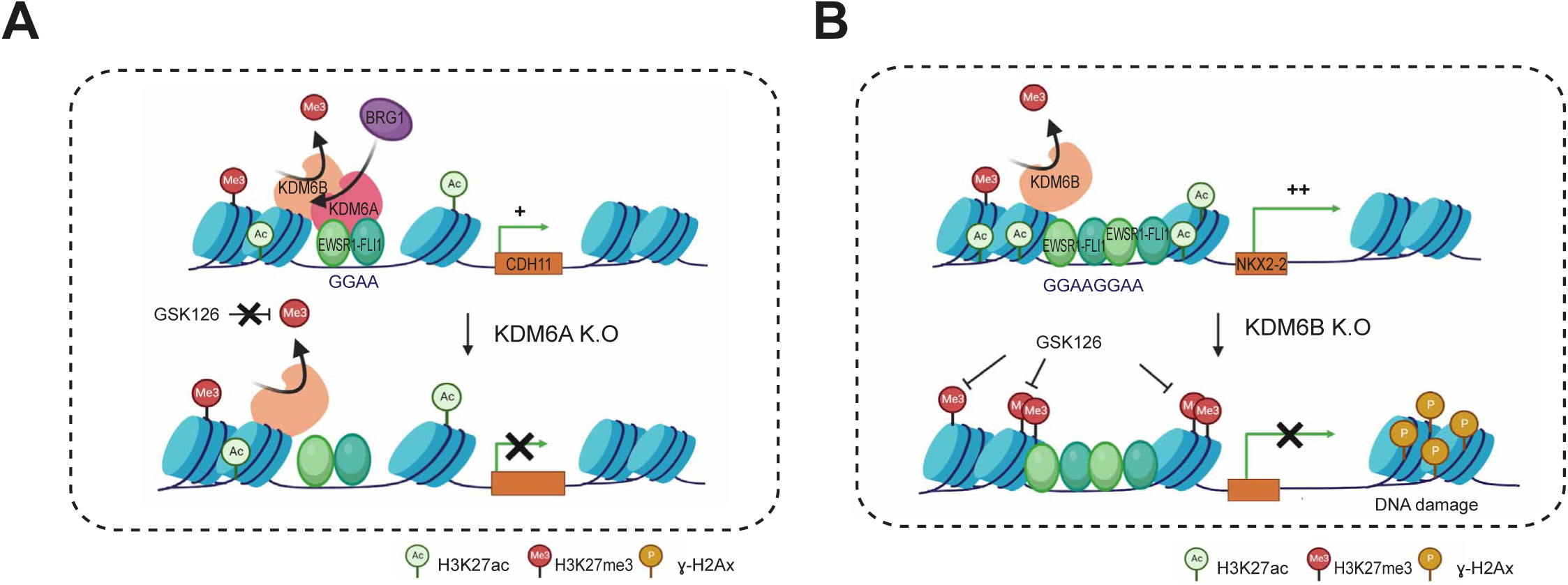
Mechanisms behind EWSR1-FLI1-mediated activation driven by KDM6A and KDM6B and the effect of its knockout. (A) KDM6A recruits BRG1 to single GGAA regions bound by EWSR1-FLI1 with or without KDM6B and low H3K27ac. Depletion of KDM6A perturbs BRG1 recruitment at these sites in a demethylase-independent manner, causing repression of target genes without changes on H3K27me3. Thus, the addition of an EZH2 inhibitor like GSK126 in KDM6A-null cells has no effect on these regions and cell viability is preserved. (B) Exclusive KDM6B-bound regions with EWSR1-FLI1 contain GGAA repeats and a strong enrichment of H3K27ac. KDM6B knockout results in the accumulation of H3K27me3 at these sites followed by repression of the target genes. Moreover, KDM6B-null cells have higher expression of double-strand DNA repair genes. Consistently, the addition of GSK126 generates a cell-stress response that in turn accentuates ɣ-H2Ax mark accumulation and cell death. Created with BioRender.com.

It has been shown that the participation of KDM6A in cancer progression is not restricted to its enzymatic activity but to a demethylase-independent function (Kim et al., 2014; Leng et al., 2020). In Ewing cells, although we observed a general increase of H3K27me3 upon KDM6A knockdown, this mark remains mostly unchanged at active EWSR1-FLI1/KDM6A-bound enhancers, suggesting a demethylase-independent function at these regions. In EwS, BRG1 and the BAF155/SMARCC1 subunits are recruited by the fusion oncogene at enhancers containing GGAA repeats to activate transcription (Boulay et al., 2017). In addition, KDM6A recruitment of BRG1 was described at cardiac-specific enhancers as a key step in activation of cardiac developmental programs (Seunghee et al., 2012). Our data reveals that KDM6A recruits the SWI/SNF member BRG1 at EWSR1-FLI1 enhancers in a demethylase-independent manner with critical consequences on transcriptional activation of these targets (Figure 7). Interestingly, while targets of enhancers containing multimeric GGAA repeats and absence of KDM6A are exclusively downregulated upon KDM6B knockout, the targets of single GGAA repeats where both demethylases co-localize are downregulated solely upon knockout of KDM6A. These results might reinforce the strong potential of KDM6A in cell identity based on its scaffolding properties of proteins such as BRG1. Indeed, only KDM6A and KDM6C contain six tetratricopeptide repeat (TPR) domains at the N terminus necessary for physical interactions with the MLL3/4 complex (Kalisz et al., 2020; Shpargel et al., 2017). However, both KDM6A and KDM6B have been implicated in condensate phase separation (Shi et al., 2021; Vicioso-Mantis et al., 2022).

Treatment with the KDM6A/B demethylase inhibitor GSK-J4 has been reported to be effective at the preclinical level for pediatric cancers like neuroblastoma and diffuse intrinsic pontine glioma (DIPG) (Hashizume et al., 2014; Lochmann et al., 2018). Here, we demonstrated that expression of EWSR1-FLI1 targets driven by KDM6B is dependent on its demethylase activity in EwS (Figure 7). Our results in KDM6B knockout cells for targets containing multimeric GGAA repeats, such as *NKX2-2* or *CCND1,* are consistent with the increase of H3K27me3 at EWSR1-FLI1 active enhancers upon GSK-J4 treatment (Heisey et al., 2021). Moreover, our results suggested that the effects of GSK-J4 observed *in vitro* and *in vivo* studies by Heisey et al. (2021), are mainly due to the specific targeting of the demethylase activity of KDM6B. Whether these observations can be extended to all EWSR1-FLI1 targets should be studied in more detail.

EZH2 sensitization upon KDM6A loss has been mostly described in cancer, including multiple myeloma, lung cancer and bladder cancer, where KDM6A acts as a tumor suppressor. In such cases, EZH2 inhibition delays tumor onset and induces tumor regression of KDM6A-null cells (Ezponda et al., 2017; Ler et al., 2017; Wu et al., 2018). Here, we observed sensitization to EZH2 inhibition in KDM6B-deleted cells (Figure 7). Nevertheless, contrary to the PRC2 amplification demonstrated in multiple myeloma and bladder cancer upon KDM6A deletion, EZH2 expression decreased upon KDM6B knockout in EwS cells. These results suggest that EwS mechanistically resembles pediatric cancers such as neuroblastoma, where both KDM6B and EZH2 are overexpressed and constitute druggable targets (Chen et al., 2018; D’Oto et al., 2021). It has been proposed that PcG proteins promote survival of EwS cells in hypoxia conditions by repression of KCNA5 (Ryland et al., 2015). Indeed, inhibition of EZH2 methyltransferase activity has no impact on cell survival under physiological conditions but results in loss of cell viability under stress conditions (i.e. hypoxia and nutrient deprivation) by induction of KCNA5. Our results suggest that KDM6B KO induces a level of stress to EwS cells, as shown by the increase in γH2AX, sufficient to activate apoptosis upon EZH2 inhibition. These results are in agreement with the specific induction of DNA repair pathways observed in KDM6B-depleted EwS cells. Besides, the non-specific demethylase inhibitor JIB-04 induces DNA damage in EwS cells (Parrish et al., 2018). Future experiments will address the differential contribution of both demethylases to DNA damage and EZH2 functionality.

## AUTHOR CONTRIBUTIONS

E.F.-B, S.S.-M. and J. M. designed and wrote the manuscript. S.S.-M. and J.M. supervised all the work conducted. E.F.-B., C. R., M.S.-J., P.T., O.M., E.P. and S.M. performed the experiments. E.B., G.F., P.C, S.P. and S.G. performed all the statistical and bioinformatic analysis. N.R., A.A., C.L., and L.dC provided expertise and feedback. All authors reviewed the manuscript.

## DECLARATION OF INTERESTS

The authors declare no competing interests.

## METHODS

### Mesenchymal stem cells and cell lines

hpMSC were extracted from the healthy bone marrow of pediatric donors of our institution and characterized according to described protocols (Riggi et al., 2008; Suva et al., 2004; Suva et al., 2008). Cells were cultured at low confluence with Iscove’s Modified Dulbeco’s Medium (Gibco) supplemented with 10% fetal newborn calf serum, 1% penicillin/streptomycin and 10 ng/mL of PDGF-BB (PeproTech).

The EwS cell lines, A673 and SK-ES-1, were purchased from ATCC. They were cultured in RPMI 1640 media (Gibco) and supplemented with 10% FBS, L-glutamine and penicillin/streptomycin. Cells were maintained in a humidified chamber at 37°C and 5% CO2, and split every 2-3 days when reaching confluence. EwS cells were treated with the EZH2 inhibitor GSK126 and the demethylase inhibitor GSKJ4 (both from Selleckchem) at 15 µM and 2.5 µM for 24 or 72 hours, respectively. DMSO was used as vehicle control.

### Patient samples

Biopsies from 45 primary EwS tumors at diagnosis from the Biobank of Hospital Infantil Sant Joan de Déu, integrated in the Spanish Biobank Network of ISCIII and in the Xarxa de Tumors de Catalunya, were used for experimental purposes in agreement with the ethical committee procedures. Two samples were not appraisable for technical reasons and were excluded.

### Lentiviral and transient transfections

Infection was performed as previously described (Sánchez-Molina et al., 2020). Infected cells were selected with 0.5-1 µg/mL puromycin for 72 hours and maintained in the first passages. Induction of the shRNA was performed with doxycycline hyclate (Sigma Aldrich) at 2 µg/mL for 72 hours. Validation of the overexpression or knockdown was determined by Western blot.

EwS cell lines were seeded in 6-well plates at 0.25·10^6^ cells/mL and transfected with Lipofectamine RNAiMAX (Invitrogene) using 30 pmol of siRNA in OptiMEM media (Gibco) following manufacturer’s instructions. Target sequences for small interfering RNA and the SMARTvector inducible Tet-On shRNA system (Dharmacon) targeting KDM6A or KDM6B are described in Table S7. Empty and EWSR1-FLI1-pLIV vector was kindly provided by Dr Nicolo Riggi and published in Boulay et al. (2017).

### CRISPR/Cas9 genome editing

KDM6A or KDM6B knockout cells were generated using the Gene Knockout Kit v2 (Synthego) containing a pool of three validated sgRNA and exogenous Cas9 following manufacturers guidelines. CRISPR-edited cell pools were isolated by limit dilution in 96-well plates. Isolated clones were expanded and cells were collected for subsequent validation by Sanger sequencing of the PCR product. The Inference of CRISPR Edits (ICE) analysis software (Conant et al., 2022) was used to analyze the obtained sequences of edited cells (clones with a model fit R^2^ > 0.6 were only considered). Knockout clones were further validated by Western blot. Target sequences for the pool of three sgRNA targeting KDM6A or KDM6B are described in Table S7.

### Cell viability and clonogenic assays

Cell viability was determined by seeding EwS cell lines at 0.25·10^5^ cells/mL in clear bottom black walled 96-well Tissue Culture plates (Thermofisher) and incubated with 10 µL of Cell Titer Blue (Promega). Fluorescence was measured using the Infinite M Nano+ (Tecan) microplate reader at 560_Ex_/590_Em_ nm. Clonogenic assays were performed by seeding 0.12·10^4^ cells/mL in 6-well plates and changing media every 2-3 days until visible colonies were grown in wells. Cells were then fixed for 10 minutes with 4% paraformaldehid (Santa Cruz Biotech), washed with PBS and then incubated with a crystal violet solution (2% W/V, 20% methanol in PBS) for 5 minutes. Quantification of the colony number was performed with the Image J (Schneider et al., 2012) pluggin ColonyArea (Guzmán et al., 2014).

### Cell cycle Analysis and Annexin V staining

Cultured cells were fixed in 70% ethanol and stained with 25 µL propidium iodide (1 mg/mL) and 25 µL RNase (10 mg/mL). Following 30 min incubation at 37°C cells were spinned by centrifugation and washed with PBS. Cell cycle analysis was performed with Novocyte Flow Cytometer (Agilent Technologies, Inc).

Cultured cells treated with the inhibitor were collected with trypsin, counted and re-suspended in Annexin binding buffer from the Alexa Fluor® 488 Annexin V/Dead Cell Apoptosis Kit (Thermofisher) and followed manufacturer’s instructions. AnnexinV-stained cells were immediately run in Cytek Aurora CS full spectrum flow cytometer (Cytek Biosciences). FlowJo software version 10.2 was used to analyze collected data.

### RNA Extraction and Real-Time Quantitative RTqPCR

Total RNA was isolated and purified using the RNeasy Mini Kit (Qiagen) following manufacturer’s instructions. Quantification of RNA samples was performed using Nanodrop 1000 spectrophotometer (Thermo Fisher Scientific). Reverse transcription (RT) was performed using 1 µg of purified RNA and converted to cDNA with the retrotranscriptase M-MLV Reverse Transcriptase, the RNasin Plus RNase inhibitor and random primers (all from Promega). SYBR Green PCR Master Mix (Applied Biosystems) was used to perform quantitative PCR (qPCR) with QuantStudio 6 Flex (Applied Biosystems) using specific primer sequences (see Supplementary Table 7). Obtained data was normalized to housekeeping gene and analyzed with the 2^-ΔΔCT^ method relative to the experimental control condition. Data was performed in at least three independent biological experiments and expressed as mean ±SEM.

### RNA-seq and Functional Analysis

RNA-seq libraries were prepared with 0,5-1 µg of total high quality RNA collected from samples and the Illumina Stranded Total RNA Prep kit (Illumina) according to manufacturer’s instructions. Fastq files were analyzed using FastQC software (Wingett and Andrews, 2018) to assess read quality. Adapters were removed and reads were trimmed using Cutadapt software (Martin, 2011) according to per base Phred quality scores and minimum length. Reads were pseudoaligned to GRCh37 using Kallisto (Bray et al., 2016). Gene read counts from Kallisto were used to determine differential gene expression using R packages tximport (Soneson et al., 2015) and DESeq2 (Love et al., 2014). ERCC spike-ins were used for sample normalization and batch effect was removed using Limma package (Ritchie et al., 2015). Heatmaps were performed with heapmap2 R package. Gene set enrichment analysis to determine the correlation between ChIP-seq targets of KDM6A and KDM6B with their respective transcriptomes was performed with GSEA (Subramanian et al., 2005) using each list of putative targets of KDM6A/KDM6B as the gene set at every running. Reports of functional enrichments of GO and other genomic libraries were generated using the EnrichR tool (Kuleshov et al., 2016). Additional RNA data was obtained from A673 shFLI1 (GSE61953), the GEO repositories including 184 EwS tumors at diagnosis (GSE17679, GSE34620 and GSE37371), and from the Cancer cell line Encyclopedia (Barretina et al., 2012).

### Immunohistochemistry

Immunohistochemical analyses were performed following standard protocols. Fixed tumor xenografts were embedded in paraffin and cut in consecutive 2 µm thick sections. Sections of paraffin tumors were deparaffinized, rehydratated in an alcohol battery and incubated with antigen retrieval with Epitope retrieval solution pH 6.0 (Novocastra Laboratories). Blocking with endogenous peroxidase was performed with Protein Block (Novocastra Laboratories) for 5 min and subsequent steps were performed in DAKO autostainer link 48 (Agilent Technologies, Inc.). Slides were counterstained in hematoxylin-eosin, dehydrated with alcohol and xylene and finally cover slipped with DPX. Nanozoomer 2.0 (Hamamatsu Photonics) was used to scan selected tumors for digital image processing. For KDM6A and KDM6B stains in EwS tumors immunohistochemical semi-quantification was scored by an independent pathologist. A semiquantitative histoscore (H-score) value was calculated based on a linear combination of the intensity and proportion of stained cells per camp that punctuates strongly stained nuclei (SSN), the percentage of moderately stained nuclei (MSN), and the percentage of weakly stained nuclei (WSN) following the following formula: H-score = 1 x WSN + 2 x MSN + 3 x SSN. The H-score value ranges possible scores from 0 to 300 (Detre et al., 1995). The list of primary antibodies and dilutions used are in Table S7.

### Protein Extract Preparation and Western blotting

Whole cell protein extracts were prepared in RIPA buffer (10 mM Tris-HCl pH 8, 1 mM EDTA, 0.5 mM EGTA, 1% Triton X-100, 0.1% sodium deoxycolate, 0.1% SDS, and 140 mM NaCl) containing phosphatase and EDTA-free Protease Inhibitor Cocktail (Roche). Cell lysates incubated on ice for 30 min and sample were centrifuged at 12,000 rpm for 15 min at 4°C. Histone extracts were isolated using the EpiQuick Histone Extraction kit (Epigentek) following manufacturer’s instructions. Protein supernatants were collected and quantified by Bradford assay (Sigma Aldrich).

50 µg and 5 µg of whole cell or histone protein extracts were mixed with loading Laemmli buffer with DTT for Western blot experiments and followed standard protocols. The primary antibodies and dilutions are listed (see Table S7) and secondary antibodies goat anti-rabbit and goat anti-mouse IRDye (Li-COR Biosciences) were diluted 1:10,000 to blotted membranes an incubated for 1 h at room temperature. Nitrocellulose membranes (Amershan Protran) were scanned and visualized with Li-COR Odyssey Infrared Imaging System (Li-COR Biosciences).

### ChIP-qPCR

ChIP-qPCR assays were performed as previously described (Sánchez-Molina et al., 2014). Cultured cells were fixed using 1% of methanol-free formaldehyde (Thermo Fisher Scientific) for 10 min at room temperature, and crosslinking was stopped by adding 500 µL of glycine (1.25 M). Lysis was performed in soft lysis buffer 0.1% SDS, 0.15 M NaCl, 1% Triton X-100, 1 mM EDTA, and 20 mM Tris pH 8) supplemented with 1 mg/mL of protease inhibitors (Roche). Cell lysates were sonicated for 10 cycles with Bioruptor Pico (Diagenode) until chromatin was sheared to an average length of 200 bp. After centrifugation, a small fraction of eluted chromatin was measured with Qubit dsDNA HS kit (Thermofisher Scientific). Immunoprecipitations were prepared starting with 30 µg for each antibody and incubated overnight at 4°C in a rotating wheel (see Table S7). 50 μL of Dynabeads Protein A (Invitrogen) were added to samples, and the slurry was incubated for 2 hours to capture DNA fragments. Immunoprecipitates were washed with the following buffers: TSE I (0.1% SDS, 1% Triton X-100, 2 mM EDTA, 20 mM Tris–HCl pH 8, 150 mM NaCl), TSE II (0.1% SDS, 1% Triton X-100, 2 mM EDTA, 20 mM Tris–HCl pH 8, 500 mM NaCl), TSEIII (0.25 M LiCl, 1% Nonidet P-40, 1% deoxycholate, 1 mM EDTA, 10 mM Tris–HCl pH 8), and Tris-EDTA buffer. All incubation and washing steps were performed in a rotating wheel at 4°C to avoid protein degradation. DNA captured by the beads was eluded by adding 120 μL of a solution containing 1% SDS, 0.1 M NaHCO3 and decrosslinked at 65°C for 3 hours with gentle shaking. Genomic DNA fragments from ChIP samples were purified with QIAquick PCR Purification kit (Qiagen) and eluted in 50-100 μL of Tris-EDTA buffer. Differences in DNA content from ChIP assays were determined by qPCR using the SQ6 Real Time PCR System and SYBR Green master mix (Applied Biosystems). The reported data from at least three independent experiments represent real-time PCR values normalized to input DNA and are expressed as percentage of bound/input signal and presented as mean±SEM.

### ChIP-seq and Bioinformatic Analysis

ChIP-seq libraries were prepared using 2-5 ng of input and ChIP samples and the kit NEBNext Ultra DNA Library Prep for Illumina (New England Biolabs) following manufacturer’s protocol. All purification steps were performed using Agen Court AMPure XP beads (Qiagen). NEBNext Multiplex oligonucleotides for Illumina (New England Biolabs) was used for library amplification. Quality control and fragment size was analyzed using Agilent High Sensitivity ChIP and quantified with KAPA Library Quantification Kit (KapaBiosystems). ChIP-seq data from DNA and input samples were sequenced with Hiseq 2500 Illumina sequencing system.

ChIP-seq samples were mapped against the hg19 human genome assembly using Bowtie with the option -m 1 to discard those reads that could not be uniquely mapped to just one region (Langmead et al., 2009). ChIP-seq samples normalized by spike-in were mapped against a synthetic genome constituted by the human and the fruit fly chromosomes (hg19 + dm3) using Bowtie with the option -m 1 to discard reads that did not map uniquely to one region. MACS was run with the default parameters but with the shift-size adjusted to 100 bp to perform the peak calling against the corresponding control sample (Zhang et al., 2008). DiffBind (Ross-Innes et al., 2012) was run next over the union of peaks from each pair of replicates of the same experiment to find the peaks significantly enriched in both replicates in comparison to the corresponding controls (DiffBind 2.0 arguments: categories = DBA_CONDITION, block = DBA_REPLICATE and method = DBA_DESEQ2_BLOCK).

We used SeqCode (Blanco et al., 2021b) for ChIP-seq downstream analysis across multiple stages: (i) the genome distribution of each set of peaks was generated by counting the number of peaks fitted on each class of region according to RefSeq annotations (O’Leary et al., 2016). Promoter is the region between 2.5 kb upstream and 2.5 kb downstream of the transcription start site (TSS). Genic regions correspond to the rest of the gene (the part that is not classified as promoter) and the rest of the genome is considered to be intergenic. Peaks that overlapped with more than one genomic feature were proportionally counted the same number of times; (ii) aggregated plots showing the average distribution of ChIP-seq reads around the TSS or along the gene body of each target gene were generated by counting the number of reads for each region according to RefSeq and then averaging the values for the total number of mapped reads of each sample and the total number of genes in the particular gene set; (iii) heatmaps displaying ChIP-seq signal strength around the summit of each peak were generated by counting the number of reads in this region for each individual peak and normalizing this value with the total number of mapped reads of the sample. Peaks on each heatmap were ranked by the logarithm of the average number of reads in the same genomic region; (iv) boxplots showing the ChIP-seq level distribution for a particular ChIP experiment on a set of genomic peaks were calculated by determining the maximum value on this region at this sample, which was assigned afterwards to the corresponding peak. To quantify genome-wide differences on H3K27me3 gain/loss, we performed a similar approach over the full set of bins of 1 kb (average value per bin, genome assembly: hg19). ChIP-seq samples containing spike-in, values were normalized for the total number of reads mapped on the fruit fly genome, as previously described (Blanco et al., 2021a); (v) BedGraph profiles were generated from each set of mapped reads and uploaded into the UCSC genome browser to generate the screenshots of tracks along the manuscript (Kent et al., 2002); (vi) the set of target genes of a biological feature was found by matching all ChIP-seq peaks in the region 2.5 kb upstream of the TSS until the end of the transcripts as annotated in RefSeq.

To build our collection of enhancers and promoters, we reanalyzed published ChIP-seq samples of H3K4me1, H3K27ac, H3K27me3, and H3K4me3 in A673 cells (Riggi et al., 2014). Each class of enhancer and promoter categories were defined as in Blanco et al. (2020). Promoters were defined as ChIP peaks of H3K27 found up to 2.5 kb from the TSS of one gene and enhancers on intergenic areas outside promoters or within gene introns. H3K4me3 was required to be present in promoters but absent in enhancers. We defined five classes of regulatory elements: active enhancers (H3K27ac), active promoters (H3K27ac + H3K4me3), poised enhancers (H3K27me3), primed enhancers (H3K27ac + H3K4me1) and bivalent promoters (H3K27me3 + H3K4me3). To construct the list of putative targets of KDM6A/KDM6B enhancers, we identified the list of genes in the vicinity of overlapping KDM6A+EWSR1-FLI1 and KDM6B+EWSR1-FLI1 modules (maximum distance between peaks and differentially regulated genes: 100 kb). Reports of functional enrichments of GO and other genomic libraries were generated using the EnrichR tool (Kuleshov et al., 2016). Motif analysis of the sequences of ChIP-seq peaks was performed with the MEME-ChIP tool, adjusting the MEME motif width between 5 and 15 bps (Machanick and Bailey, 2011). ChIP-seq raw data from H3K27me3 in control and EWSR1-FLI1 hpMSC was kindly provided by Dr Nicolo Riggi and Dr Miguel Rivera (accession number GSE106925). For ChIP-seq experiments with EWSR1-FLI1 in EWS cell lines we used a new antibody (see Supplementary Table 7). We proved that at peak level it correlates 62% with published data (Riggi et al., 2014) and nicely reproduce our previous data with RING1B overlap (Sánchez-Molina et al., 2020).

### Murine xenograft studies

*In vivo* studies were performed after the approval of the Institutional Animal Research Ethics Committee and by the animal experimentation commission of the Catalan government. 1·10^6^ cells of parental, KDM6A knockout (sgRNA#1 and 2) and non-targeting control (sgCTRL) A673 cells in 200 µL of PBS with Matrigel (Becton Dickinson) were subcutaneously injected in two flanks of Athymic Nude mice (Envigo) (n=11 for A673, sgKDM6A#1 and 2; n=9 for sgCTRL). Tumor growth was monitored three times per week by measuring growing tumors with a digital caliper. Mice were sacrificed when tumors reached a volume of 300 mm^3^ and tumors were excised. Collected tumors were divided in parts: one part was frozen for protein experiments and the other was fixed in 10% formalin for immunohistochemistry experiments. For RNA experiments 4 tumors of each experimental group were dissociated with collagenase IV (50 mg/mL) (Sigma) and a tissue chopper (Ted Pella, Inc). Then tissue homogenates were digested with 5 mg/mL of DNAse I (Sigma) and trypsin/EDTA (0.25%) to subsequently separate mouse stroma cells from human cells with Mouse Cell Depletion kit (Milteny Biotech) following manufacturer’s guidelines. Log-rank test was used to calculate significance of the groups in Kaplan-Meier.

### Quantification and statistical analyses

Data was analyzed using Graphpad Prism 9 software (San Diego, CA, USA) version v.9.1.2 and expressed as mean ±SEM or SD as indicated in figure legends. Kruskal-Wallis one-way analysis of variance (ANOVA) and two-way ANOVA with Tukey’s correction (for non-normally distributed data) were applied to determine differences between multiple groups. Student t-test and Mann-Whitney t-test (for non-normally distributed data) were used for non-paired comparisons of two groups. Kaplan-meier curves were compared with the log-rank (Mantel-Cox) test. *P<0.05, **P<0.01, ***P<0.001.

### Data accession numbers

Raw data and processed information of the ChIP-seq and RNA-seq experiments generated in this article were deposited in the National Center for Biotechnology Information Gene Expression Omnibus (NCBI GEO) (Barrett et al., 2012) repository under the accession number GSE211743.

## ACKNOWLEDGEMENTS

We thank M. Martínez-Balbás for critical reading of the manuscript. We also thank M. Suñol and Neus Prats for technical advice. Last, we are grateful to the Band of Parents at Hospital Sant Joan de Déu for supporting the overall research activities of the Cancer Pediatrics Group (IRSJD and PCCB). E.F.-B. was supported by the Spanish government grant, Instituto de Salud Carlos III (PI16/00245) to J.M. The work in the L.D.C. laboratory is supported by grants from the Spanish of Economy, Industry and Competitiveness (MEIC) (PID2019-108322GB-100) and from AGAUR.

## Notes

### Competing Interest Statement

The authors have declared no competing interest.

