## Supplementary Figures for "KDM6 demethylases mediate EWSR1-FLI1-driven oncogenic transformation in Ewing Sarcoma"

**SUPPLEMENTARY INFORMATION**

**Figure S1. H3K27me3 genome-wide redistribution upon EWSR1-FLI1 overexpression in hpMSCs, Related to Figure 1.**

(A) Metagene plot showing H3K27me3 (left) and H3K27ac (right) ChIP-seq signals in 6,150 and 15,810 target genes, respectively, at transcription start site (TSS) and transcription end site (TES) within 5000 kb window in control (CTRL) and upon infection with the oncogene (EWSR1-FLI1) in hpMSC.

(B) Scatter plot of H3K27me3 signal in 3,069,655 bins of 1 kb in CTRL (x-axis) and EWSR1-FLI1 (y-axis) hpMSC ($R^2$=0.582, slope=0.703). Bins that significantly gain (Up bins) or loose H3K27me3 (Down bins) are highlighted in red or blue, respectively.

(C) Bar plot depicting percentage of annotated regulatory elements (active/poised/primed enhancers and active/poised promoters) covered by at least one bin of the genome (1 kb) in up bins or down bins of H3K27me3 in CTRL and EWSR1-FLI1 hpMSC.

(D) Boxplot depicting the average ChIP-seq signal of H3K27me3 in Up and Down bins in CTRL and EWSR1-FLI1 hpMSC.

(E) Bar chart representing the top five enriched signaling pathways of 2879 genes from Down bins (above) and 2621 genes from Up bins (below) from CTRL and EWSR1-FLI1 hpMSC and their associated P-value.

(F) Boxplot depicting immunohistochemical score (H-score) mean values for KDM6A and KDM6B in our cohort of 45 EwS primary tumors (right). Individual H-score values are represented as dots. Table shows descriptive statistics information of the recruited samples (left).

(G) Scatter plot of individual H-score values for KDM6A (x-axis) and KDM6B (y-axis) (r=0.000, CI=(-0.308,0.308), P-value=1.00).

Error bars indicate SD (D and F).

**Figure S2. KDM6A and KDM6B co-localize genome-wide with EWSR1-FLI1 at primed and active enhancers, Related to Figure 2.**

(A) Volcano plot of the significant peaks identified by DiffBind for KDM6A, KDM6B, and EWSR1-FLI1 (3737, 2687, and 4800 respectively) (P value $\leq$ 0.05 for KDM6A and EWSR1-FLI1, and FDR < $10^{-5}$ for KDM6B) in A673 cells.

(B) Table showing the top MEME DNA motifs and the corresponding E-value for every set of peaks.

(C) Bar chart representing the top five enriched gene ontology (GO) biological process of the 1511 or 1205 genes associated to KDM6A (above) or KDM6B peaks (below), respectively, and their associated P-value.

(D) Venn diagram showing the overlap between KDM6A, KDM6B, and EWSR1-FLI1 at peak level in SK-ES-1 cells.

(E) Boxplot depicting the average ChIP-seq signal of H3K4me3 (left) and H3K27me3 (right) in each set of peaks in A673 cells.

(F) Same as (E) showing the average ChIP-seq signal of RING1B in each group of peaks in A673 cells.
(G) UCSC genome browser signal tracks for KDM6A, KDM6B, EWSR1-FLI1, H3K27ac, and H3K4me1 at the *SMYD3* gene in A673. EWSR1-FLI1 and KDM6A peaks with or without KDM6B (A-B-EF or A-EF, respectively) are represented as black bars below tracks.
(H) ChIP-qPCR of KDM6A and KDM6B in a set of EWSR1-FLI1-bound enhancer regions with both KDM6A and KDM6B (A-B-EF) or only KDM6B (B-EF) in A673 cells. *ENC1* was used as a negative control region. Error bars indicate SD (E and F). Error bars in (H) indicate SEM of three independent biological experiments

**Figure S3. Knockdown of KDM6A and KDM6B downregulates EWSR1-FLI1-activated targets, Related to Figure 3.**
(A) Western blot showing levels of KDM6A and KDM6B in whole cell extracts upon KDM6A (above) or KDM6B (below) knockdown with two shRNA sequences (sh#1 and sh#2) at 72 hours in SK-ES-1 cells. Tubulin was used as loading control. Numbers below represent band quantification of KDM6A or KDM6B normalized to tubulin and relative to shCTRL.
(B) Donut charts representing the percentage of significantly upregulated or downregulated targets upon KDM6A (above) or KDM6B (below) knockdown with sh#1 and sh#2 in A673 cells (492 and 502 genes for shKDM6A#1 and #2, respectively; 1090 and 1369 genes for shKDM6B#1 and #2, respectively).
(C) Same as (B) in SK-ES-1 cells (583 and 731 genes for shKDM6A#1 and #2, respectively; 2975 and 196 genes for shKDM6B#1 and #2, respectively).
(D) Bar chart representing the top five enriched gene ontology (GO) biological process of the 64 genes in the vicinity of KDM6A-EWSR1-FLI1 peaks (above) and the 139 genes in the vicinity of KDM6B-EWSR1-FLI1 peaks (100 kb) (below) that are significantly downregulated upon knockdown of each demethylase and their associated P-value.
(E) RT-qPCR determination of mRNA expression of EWSR1-FLI1 targets with active enhancers in shCTRL and shKDM6A (#1 and 2) in A673 cells. Values were normalized to *GAPDH* and relative to shCTRL.
(F) Same analysis as in (E) for shCTRL and shKDM6B (#1 and 2) in A673 cells.
Statistical significance was determined by Kruskal-Wallis one-way ANOVA test. Error bars indicate SEM (E and F) of four independent biological experiments; $^{**}P < 0.01$, and $^{*}P < 0.05$.

**Figure S4. KDM6A recruits BRG1 to EWSR1-FLI1-activated enhancers in a demethylase-independent manner, Related to Figure 4.**
(A) Western blot showing levels of H3K27me3, H3K27ac, and H3K4me1 in histone extracts upon KDM6A knockdown with a Tet-On shRNA doxycycline-inducible system with two shRNA sequences (#1 and #2) at 72 hours in A673 (left) and SK-ES-1 (right) cells. Histone H3 was used as loading control. Numbers below represent band quantification of H3K27me3 normalized to H3 and relative to shCTRL.

(B) Pie chart showing genomic distribution of H3K27me3 peaks relative to functional categories including promoter (±2.5kb from TSS), gene body (intragenic region not overlapping with promoter) and intergenic (rest of the genome) in control (sgCTRL) and sgKDM6A knockout (sgRNA#1) in A673 cells.
(C) Metagene plot showing H3K27me3 ChIP-seq signal of 161 KDM6A-repressed targets from RNA-seq data at transcription start site (TSS) within 5000 kb window in sgCTRL and sgKDM6A#1.
(D) Boxplot depicting the average ChIP-seq signal of H3K27me3 in 1 kb bins in sgCTRL and sgKDM6A#1. Bin mapping analysis identified 881,014 and 88,914 bins that gained (up bins) or loss (down bins) H3K27me3 signal, respectively, upon KDM6A knockout compared to control.
(E) Scatter plot of H3K27me3 ChIP-seq signal in 3,069,655 bins of 1 kb in sgCTRL (x-axis) and sgKDM6A#1 (y-axis) ($R^2$=0.272, slope=0.748). Bins that significantly gain or lose H3K27me3 are highlighted in red or blue, respectively.
(F) Venn diagram depicting the overlap between the 11658 genes associated to the bins that gain H3K27me3 upon knockout (up bins) with the 341 targets activated by KDM6A from RNA-seq data.
(G) ChIP-qPCR of H3K27me3 enrichment in the enhancer region of *TSPAN13, SYT1* and *IRS2* genes upon KDM6A knockdown with shRNA#2 in A673 cells. *DICER* and *TAL1* were used as negative and positive control regions, respectively.
(H) Same as (F) showing enrichment of BRG1 upon KDM6A knockdown with shRNA#2.
Error bars indicate SD (D) or SEM (G and H) of three independent biological experiments.

**Figure S5. KDM6A is a critical factor for EwS engraftment and tumor growth, Related to Figure 5.**
(A) Bar charts showing number of colonies from Fig. 5B, in parental, sgCTRL and sgKDM6A (sgRNA #1 and 2) A673 cells.
(B) Aggregated bar plot representing the percentage of cells on each cell cycle phase in sgCTRL and sgKDM6A (sgRNA #1 and 2) in A673 cells.
(C) Spaghetti plots showing tumor volume of xenograft tumors from parental, sgCTRL, and sgKDM6A (sgRNA #1 and 2) in A673 cells within 40 days post-injection.
(D) Boxplot representing tumor volume at 17 days post-injection of parental, sgCTRL, and KDM6A knockout cells (sgRNA#1 and #2) in A673. Each dot represents an individual tumor volume.
(E) Western blot showing levels of KDM6A in 4 representative xenograft tumors derived from sgCTRL and sgKDM6A (sgRNA#1 and 2) in A673. Tubulin was used as loading control.
(F) Boxplot representing mRNA levels of *NEFH* in a panel of 22 EwS cell lines extracted from Barretina et al. (2012). Each dot represents individual values for a given cell line with A673 cell line colored in red.
(G) Boxplot representing mRNA levels of *NEFH* in primary tumors from GEO public data repositories including EwS among other primary sarcoma tumors including

osteosarcoma (OS), rhabdomyosarcoma (RMS), and synovial sarcoma (SS). MSCs derived from the healthy bone marrow were included as control cell lines.
Statistical significance was determined by Kruskal-Wallis one-way ANOVA test (A and D) with Dunn's multiple comparison test (D). Error bars in (A, B, D, F and G) indicate SEM; $**P < 0.01$.

**Figure S6. KDM6B knockout sensitizes EwS cells to the EZH2 inhibitor GSK126, Related to Figure 6.**
(A) Western blot showing levels of H3K27me3, H3K27ac, and H3K4me1 in histone extracts upon KDM6B knockdown with a Tet-On shRNA doxycycline-inducible system at 72 hours in A673 and SK-ES-1 cells. Histone H3 was used as loading control. Numbers below represent band quantification of H3K27me3 normalized to H3 and relative to shCTRL.
(B) ChIP-qPCR of H3K27me3 enrichment in a set of KDM6B-bound targets with KDM6A at the enhancer region of *CDH11* and *IRS2* genes upon KDM6B knockout with sgRNA#2.
(C) Western blot showing levels of H3K27me3 in histone extracts of SK-ES-1 cells treated with vehicle or the demethylase inhibitor GSKJ4 at 2.5 and 5 µM (+ and ++, respectively) for 72h. Histone H4 was used as loading control. Numbers below represent band quantification of H3K27me3 normalized to H4 and relative to vehicle control.
(D) RT-qPCR determination of EWSR1-FLI1 targets with both KDM6A and KDM6B or with KDM6B ChIP-seq peaks (A-B-EF and B-EF groups, respectively) in SK-ES-1 cells treated with vehicle or GSK-J4 at 2.5 µM for 72 hours. *TBP* was used as housekeeping gene.
(E) Bar charts representing colony number, from colonies in Figure 6E, in parental, sgCTRL and sgKDM6A (sgRNA #1 and 2) A673 cells.
(F) Western blot showing levels of KDM6A, cleaved-PARP-1 (c-PARP), and phosphorylation of S139 in variant gamma-H2A.x (ɣ-H2Ax) in whole cell extracts (above) and H3K27me3 in histone extracts (below) from sgCTRL and sgKDM6A#1 A673 cells treated with vehicle or GSK126 inhibitor (15 µM for 24 hours). Tubulin and histone H3 were used as loading controls for whole cell or histone extracts, respectively. Numbers below represent band quantification of H3K27me3 normalized to H3 and relative to sgCTRL or sgKDM6A#1 treated with vehicle.
(G) Annexin V staining of sgCTRL and sgKDM6A#1 cells treated with vehicle or GSK126 inhibitor at 15 µM for 24 hours. Numbers indicate percentage of cells in each quadrant.
(H) Western blot showing levels of H3K27me3 in histone extracts of SK-ES-1 cells treated with vehicle, 10 µM of GSK126, 2.5 µM of GSKJ4 or combination of both GSK126 and GSKJ4 for 72 hours.
(I) Aggregated bar plot representing percentage of viable, early apoptosis, late apoptosis or necrotic cells from Annexin V staining upon treatment with vehicle, single and combination treatments of GSK126 and GSKJ4 after 72 hours in SK-ES-1 cells. Red brackets indicate early apoptosis statistic comparisons between experimental groups.

Statistical significance was determined by Kruskal-Wallis one-way ANOVA test (E), Mann Whitney test (D) and two-way ANOVA followed by Tukey's post hoc test. Error bars in (B, D, E and I) indicate SEM of three independent biological experiments; ***$P < 0.001$ and **$P < 0.01$.

#### SUPPLEMENTARY TABLE LEGENDS

**Table S1.** Excel file showing genes associated to gain or loss of bins of H3K27me3 (up or down bins, respectively) following EWSR1-FLI1 introduction in hpMSC.

**Table S2.** Excel file with summary of KDM6A, KDM6B and FLI1 Diffbind peaks from ChIP-seq data in A673 cells.

**Table S3.** Excel file with the associated list of genes for KDM6A and KDM6B ChIP-seq peaks in A673 cells.

**Table S4.** Excel file showing differential expressed genes of shKDM6A and shKDM6B with shRNA sequences #1 and #2 in A673 cells and the core enriched EMT gene sets upon knockdown of KDM6A or KDM6B.

**Table S5.** Excel file with the list of EWSR1-FLI1-KDM6A or EWSR1-FLI1-KDM6B targets downregulated upon KDM6A or KDM6B knockdown in A673 cells.

**Table S6.** Excel file with the list of genes from the Double Strand Break Repair category from GSEA analysis significantly upregulated upon KDM6B knockdown in A673 cells.

**Table S7.** Excel file with information of the antibodies, primers, and RNAi and CRISPR sequences used.

### Supplementary Figure 1

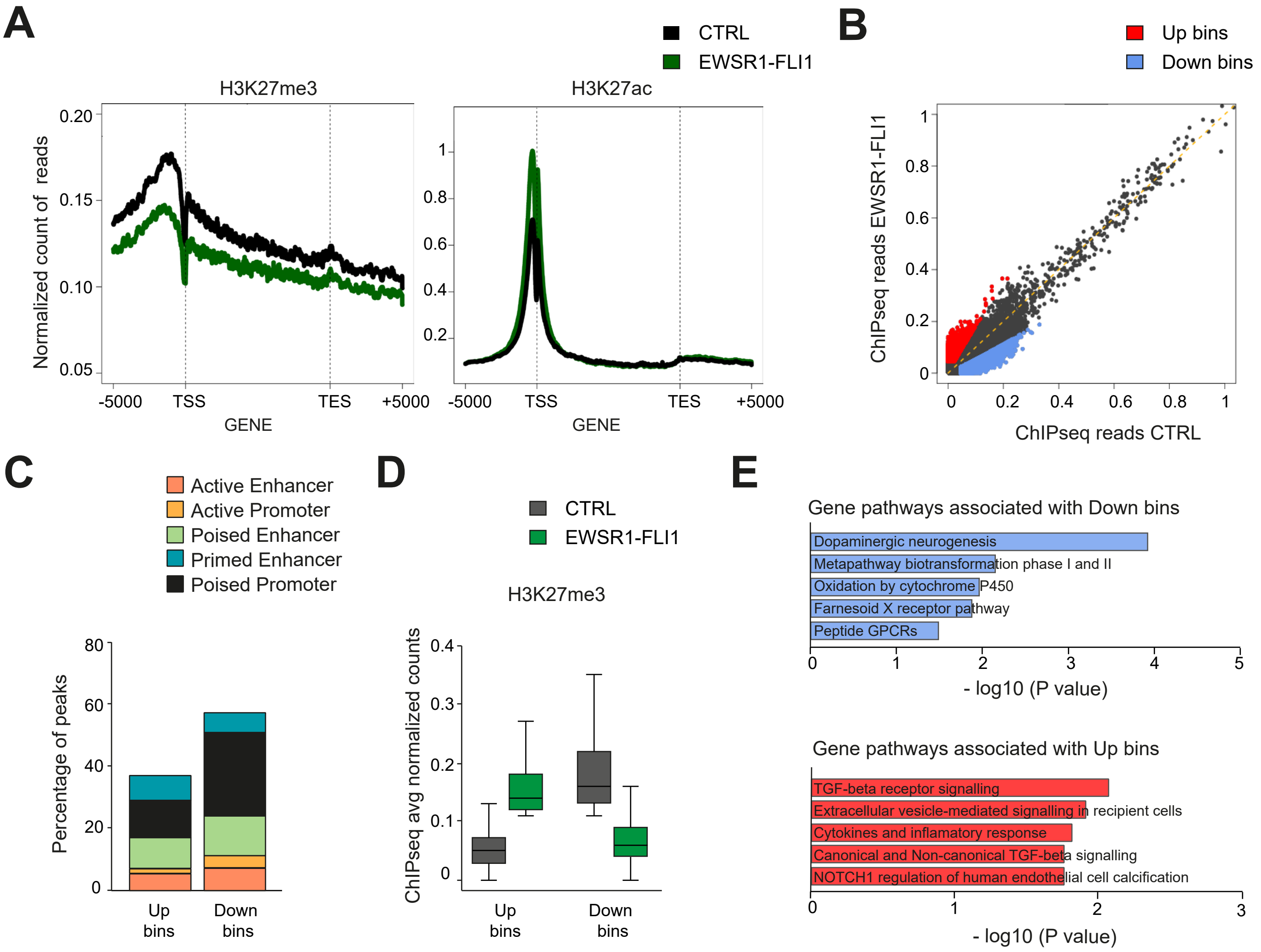


F

| | $n = 45$ | $n$ |
|---|---|---|
| H-score KDM6A | 200 [100;300] | 43 |
| H-score KDM6B | 210 [100;300] | 43 |
| Gender: | | 45 |
| Female | 16 (35.6%) | |
| Male | 29 (64.4%) | |

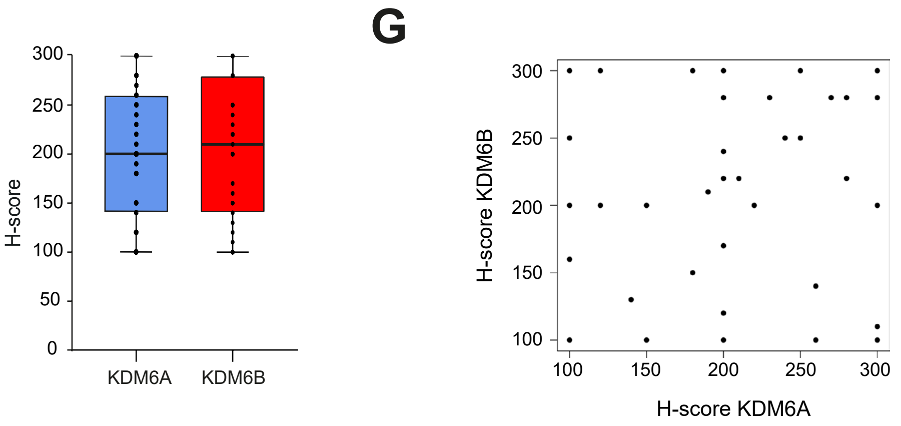

### Supplementary Figure 2

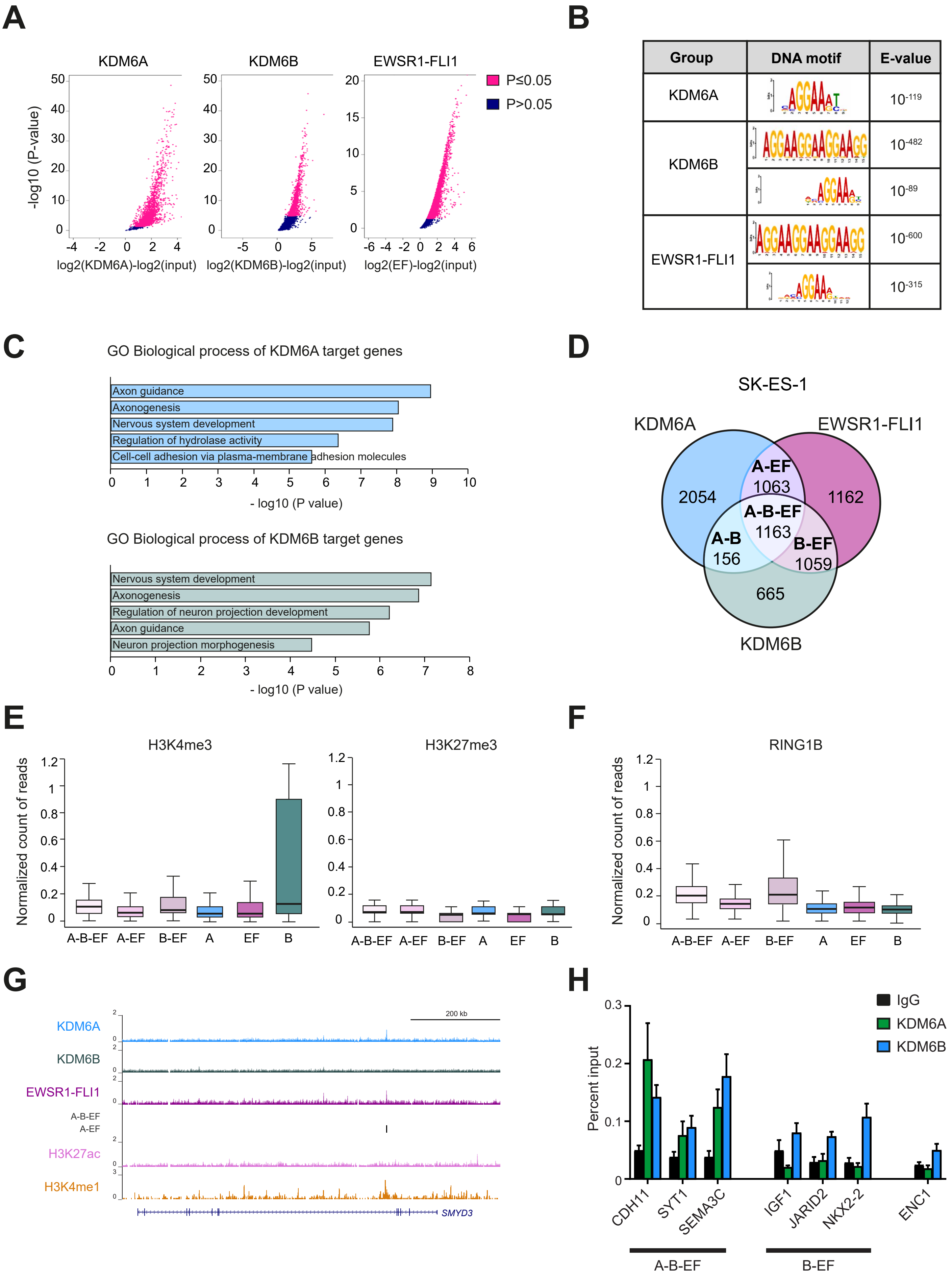

### Supplementary Figure 3

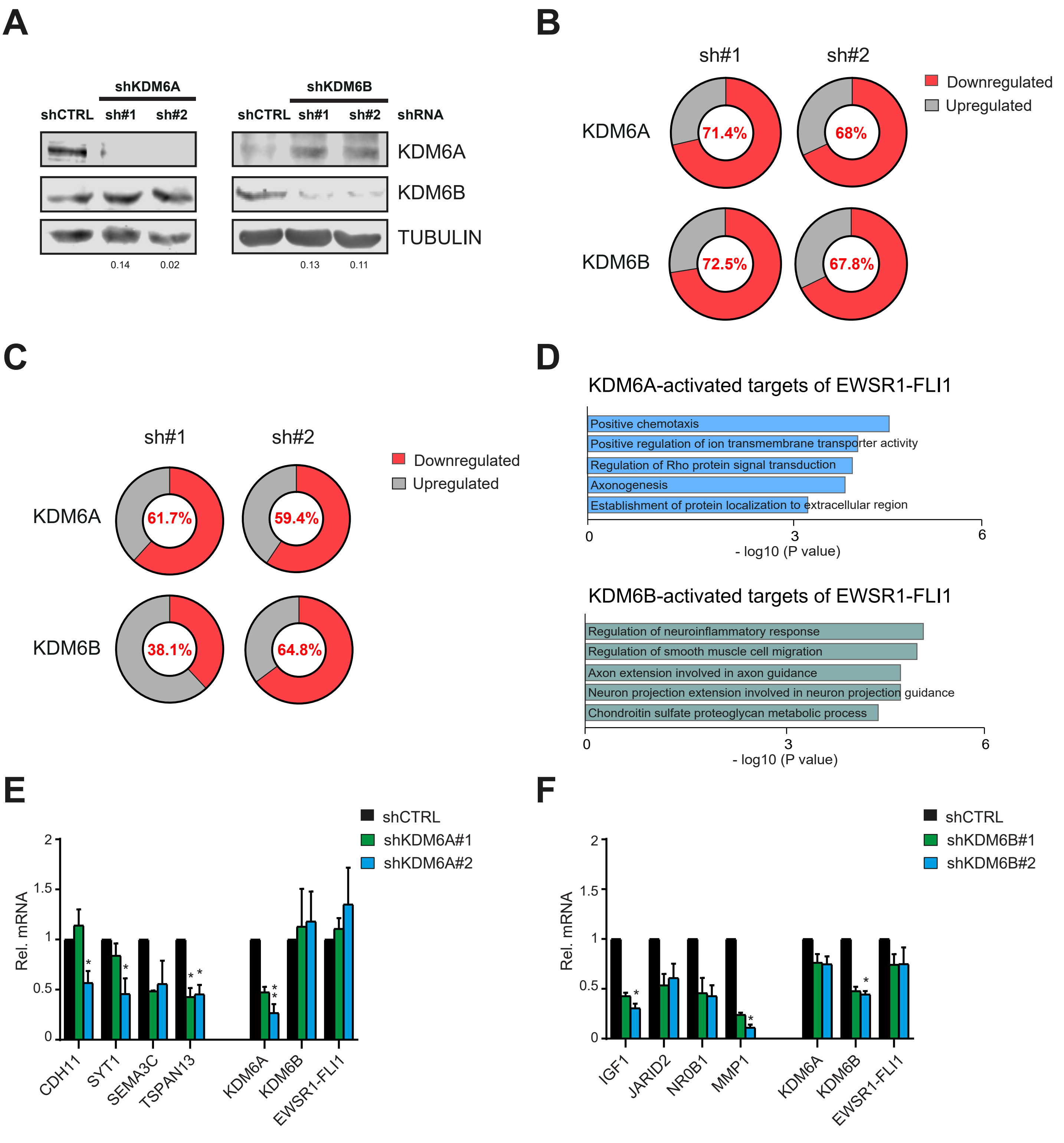

### Supplementary Figure 4

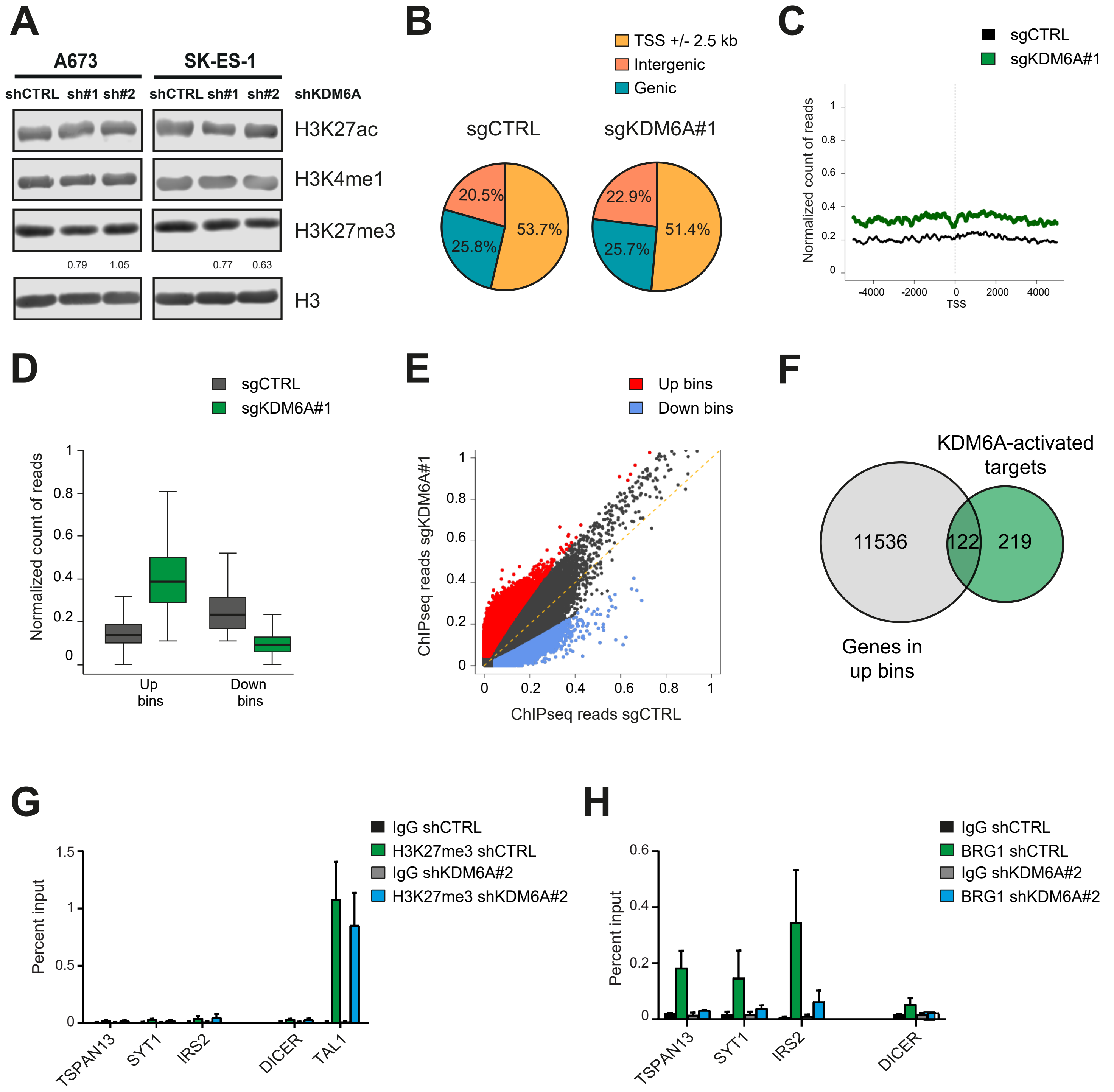

### Supplementary Figure 5

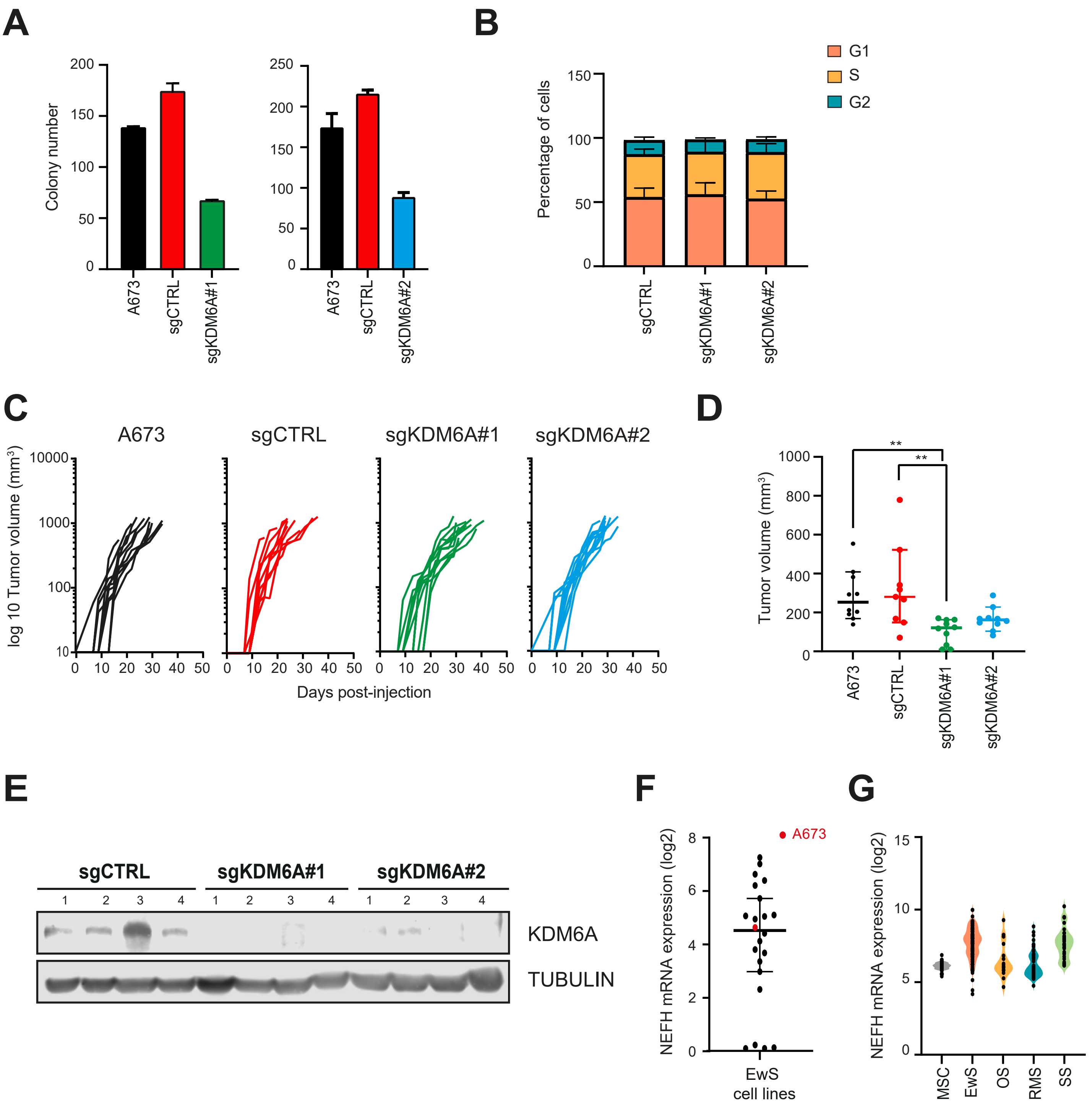

### Supplementary Figure 6

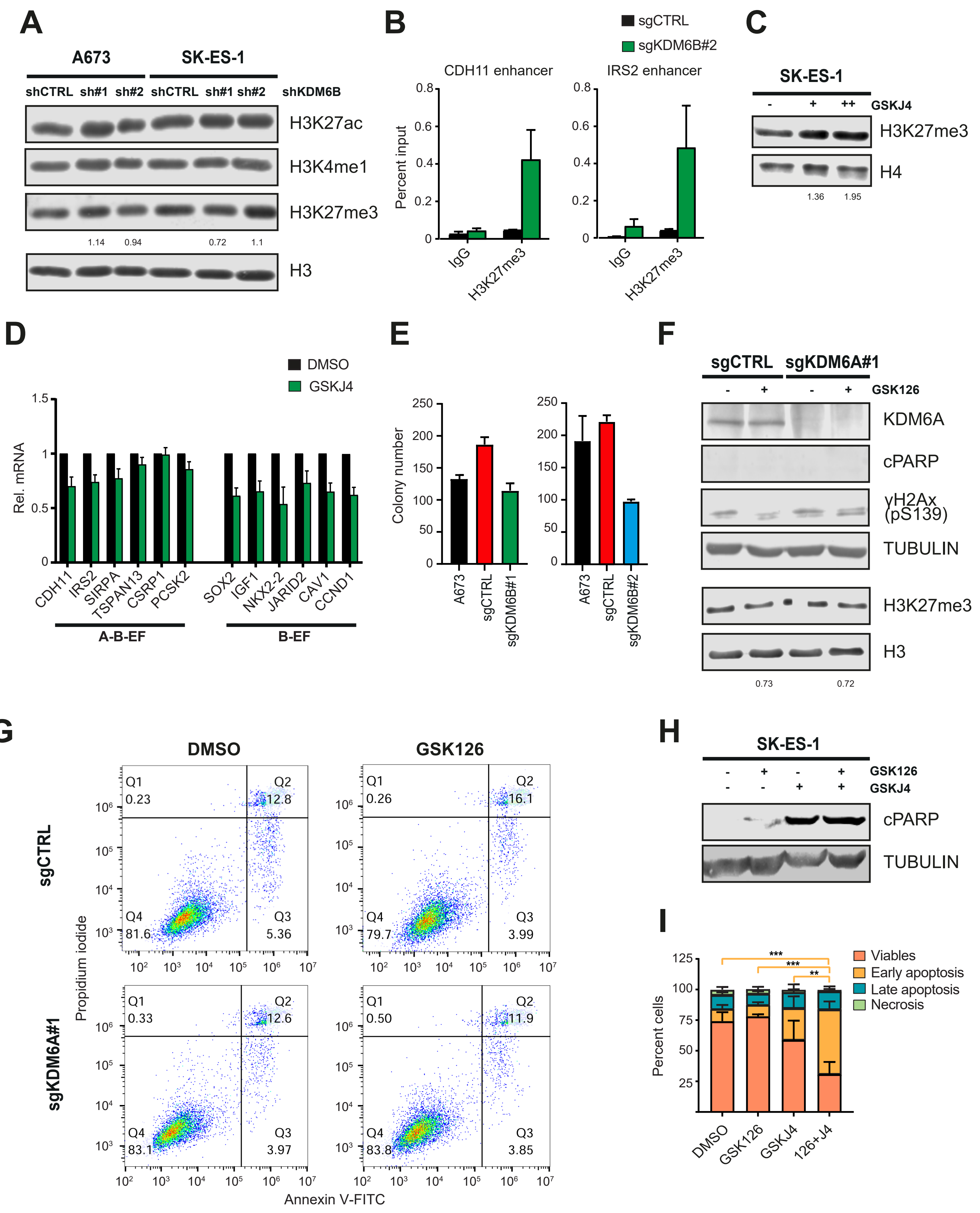
